# A universal molecular mechanism driving aging

**DOI:** 10.1101/2024.01.06.574476

**Authors:** Wan Jin, Jing Zheng, Yu Xiao, Lingao Ju, Fangjin Chen, Jie Fu, Hui Jiang, Yi Zhang

**Affiliations:** Department of Biological Repositories, Human Genetic Resources Preservation Center of Hubei Province, Zhongnan Hospital of Wuhan University, Wuhan, China; Euler Technology, ZGC Life Sciences Park, Beijing, China; National Institute of Biological Sciences, Beijing, China; Wuhan Research Center, Chinese Academy of Medical Sciences, Wuhan, China; High Performance Computing Center, the Peking-Tsinghua College of Life Sciences, Peking University, Beijing, China; Hongkong University of Science and Technology, Hong Kong, China

**Keywords:** Aging, epigenetics, DNA G-quadruplex (G4), mitosis, progeroid

## Abstract

How cell replication ultimately results in aging and the Hayflick limit are not fully understood. Here we show that clock-like accumulation of DNA G-quadruplexes (G4s) throughout cell replication drives conserved aging mechanisms. G4 stimulates transcription-replication interactions to delay genome replication and impairs DNA re-methylation and histone modification recovery, leading to loss of heterochromatin. This creates a more permissive local environment for G4 formation in subsequent generations. As a result, G4s gradually accumulate on promoters throughout mitosis, driving clock-like DNA hypomethylation and chromatin opening. In patients and *in vitro* models, loss-of-function mutations in the G4-resolving enzymes WRN, BLM and ERCC8 accelerate the erosion of the epigenomic landscape around G4. G4-driven epigenomic aging is strongly correlated with biological age and is conserved in yeast, nematodes, insects, fish, rodents, and humans. Our results revealed a universal molecular mechanism of aging and provided mechanistic insight into how G-quadruplex processor mutations drive premature aging.

## Introduction

Aging is a fundamental property of life. In his seminal study, Hayflick^1^ co-cultured cells of different *in vitro* passage numbers. By tracking their clonal progenies by karyotype counting, Hayflick showed that aged cells produce fewer progenies and are eventually outcompeted from the population. Quantitative analysis of these experiments led to his proposal of the so-called Hayflick limit, namely, that every cell could undergo only finite numbers of mitoses. Since then, modern studies have identified multiple characteristic molecular hallmarks of aged cells^2^. The cell-autonomous features of aging include genomic instability^3,4^, telomeric attrition^5^, epigenomic landscape erosion^6–11^, transcriptional changes^12–14^, proteostasis imbalance^15–17^, and metabolic switches^18^. However, the exact molecular mechanism driving aging across mitosis leading to the Hayflick limit has not been determined.

While some of the molecular processes that occur during aging, such as DNA damage, are irreversible, aged somatic cells can be “rejuvenated” back to a young ground state by nuclear transfer^19,20^, indicating that the genetic composition of aged cells is insufficient for determining the aged state. Forced expression of embryonic stem cell-specific transcription factors (TFs) in somatic cells reprograms them back to an induced pluripotent state^21^, not only reversing their differentiation state but also completely resetting their “epigenetic age” ^7,8^, resulting in an embryonic state that could rederive a young individual animal^21^. These results indicate that age is a property that is carried by non-genetic components in the cell. In agreement with this observation, chemical compounds inducing epigenetic changes could induce pluripotent reprogramming from somatic cells^22–24^. Conversely, programmed attrition of epigenetic modifier enzymes by genetic manipulation results in accelerated aging of animals^25^. Hence, aging is likely to arise from gradual erosion of the established epigenetic landscape.

Research on human progeroid syndromes (premature aging) has provided insight into the genetic mechanism driving aging. Progeroid syndromes are caused by mutations in three classes of genes. The first class of genes underlying syndromes, such as Hutchinson-Gilford progeria syndrome (LMNA^26–29^) and Marbach-Rustad progeroid syndrome (LEMD2^30^), encode proteins located in the nuclear membrane that are essential for heterochromatin anchoring. Mutations in these genes result in non-specific disruption of heterochromatin structure. The second class of genes, which cause diseases such as Fontaine progeroid syndrome, are essential for maintaining metabolic activity and proliferation, such as the mitochondria phosphate carrier SLC25A24^31,32^.

The last group of genes causes progeroid disease and includes genes associate with XFE syndrome (ERCC4^33,34^), RECON progeroid syndrome (RECQL^35^), Werner syndrome (WRN/RECQL2^4,36–39^), Bloom syndrome (BLM/RECQL3^40^), and Cockayne syndrome (CSB/ERCC6^41^ and CSA/ERCC8^34,42^). Although all of these genes function in the DNA repair machinery, they have an additional common feature: they are responsible for the resolution of non-B DNA structures, particularly G-quadruplexes (G4s) ^43–46^. G4s are single-stranded DNA structures that can spontaneously form from single-stranded DNA in open chromatin^47–53^. It is particularly enriched at transcription start sites (TSSs) ^54,55^ and at DNA replication origins^54,56,57^ and might serve as a core promoter for both transcription and DNA replication. The G4 loci are enriched in structural variation hotspots in cancer^58–61^, suggesting that these loci are particularly vulnerable to DNA breaks. This effect of G4 on DNA damage might be attributed to its ability to induce transcription-replication interactions and inhibit DNA replication^45,55,59,61–66^. Furthermore, the formation of G4s regulates chromatin accessibility^55,56,67^ and DNA methylation^68^. In addition to its function in promoting DNA damage, how G4s are linked to aging has not been determined.

Progeroid syndromes resulting from mutations in G4 resolving enzymes, especially ERCC6 and ERCC8, are phenotypically very severe and onset at an early age. Fibroblasts carrying G4 resolving enzyme mutation show accelerated decline of replicative potential *in vitro*^69^, which could not be solely explained by telomere shortening^70^ or increased DNA damage^71^. Aged cells from progeroid syndrome patients carrying WRN^72^, BLM^73^, or ERCC6^74–76^ mutations could still be “rejuvenated” back to iPSC. These data suggest that G4 may mediate non-genetic mechanisms underlying aging.

## Results

### Clock-like formation of G4 foci during cell aging

We analyzed the cellular content of G4s during aging in *in vitro* passaged primary mouse fibroblasts (pMEFs) using scFv-G4-antibody (BG4)-mediated immunofluorescence (Fig. 1a). As previously reported^48^, BG4 stained clustered G4 spots within the nuclei (Fig. 1d). We found that the number of G4 spots per nucleus was significantly elevated during cell passage (Fig. 1d-e). We then performed a quantitative BG4 CUT&Tag^51^ assay followed by qPCR and sequencing to measure G4 across the genome. We found that G4s tend to gradually accumulate in accessible chromatin (Extended Data Fig. 1), and the fraction of G4 reads within the aging-associated ATAC peak, determined as the ATAC peaks that gradually open during aging^7^, linearly increases as the cell number increases (Fig. 1b). Furthermore, the coefficient of variation of the G4 signal increased with cell age (Fig. 1c), indicating that the heterogeneity of G4 decreased during aging. These results indicate that G4s accumulate at specific genomic loci during aging.

**Figure 1.**
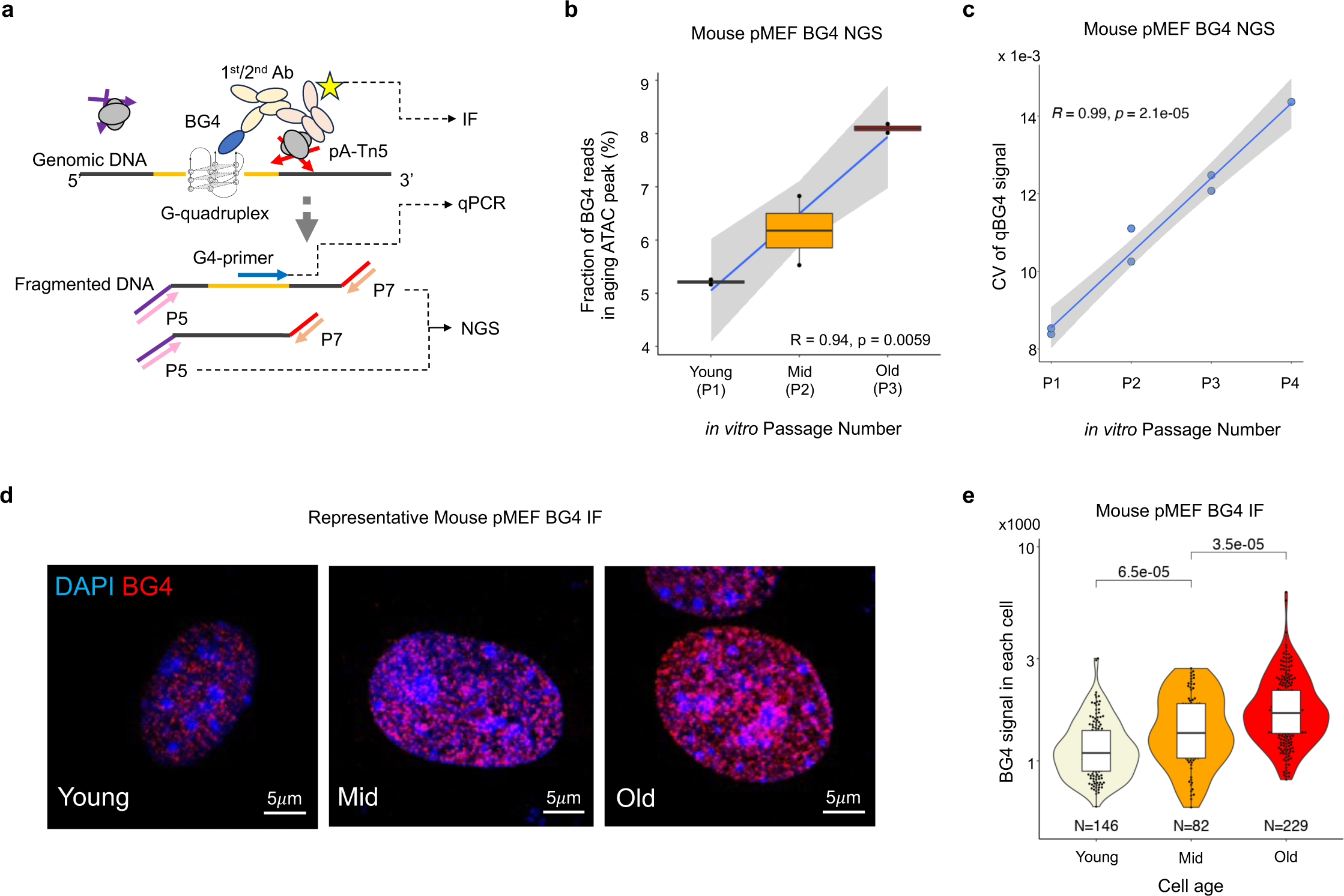
Clock-like formation of DNA G-quadruplex foci during aging. (a) Schematic of the analysis. DNA G-quadruplexes in mouse primary embryonic fibroblasts (pMEFs) were captured by a BG4 single-chain antibody and subjected to either immunofluorescence (IF) staining or pA-Tn5-mediated CUT&Tag fragmentation followed by qPCR or next-generation sequencing (NGS). **(b)** Fraction of BG4 CUT&Tag reads falling in aging-associated ATAC peaks. **(c)** Coefficient of variation of loci-wise BG4 qCUT&Tag signals in pMEF samples at different *in vitro* passages. **(d)** Representative images of BG4 immunofluorescence staining in young, middle-aged or old pMEFs. **(e)** Quantitative measurement of the BG4 immunofluorescence signal in individual pMEFs. *p*-value: t-test.

### G4 accumulation drives clock-like chromatin opening

G4s can only form on opened chromatin. Using chromatin accessibility sequencing (ATAC-seq), we found that at certain genomic loci, chromatin accessibility gradually increases during *in vitro* passage through pMEFs. Chromatin opening during aging is much stronger at TSS of genes harboring a putative G-quadruplex sequence (pGQS) in their promoters (hereinafter termed “pGQSpos genes” for simplicity) than in those without pGQS in their promoters (hereinafter termed “pGQSneg genes”) (Fig. 2a). We calculated aging-associated chromatin opening kinetics as the Z-scaled correlation coefficient between ATAC peak reads and sample age^7^ (hereinafter termed the “aging coefficient” for simplicity). The genes that showed a significant correlation between TSS chromatin accessibility and sample age (defined as those with an aging coefficient >= 3) were termed “aging genes”, which showed a significantly greater gain of chromatin opening during aging (Fig. 2a). Interestingly, such aging-associated chromatin opening is associated with concurrent gain of G4 signals at the same promoters (Fig. 2b).

**Figure 2.**
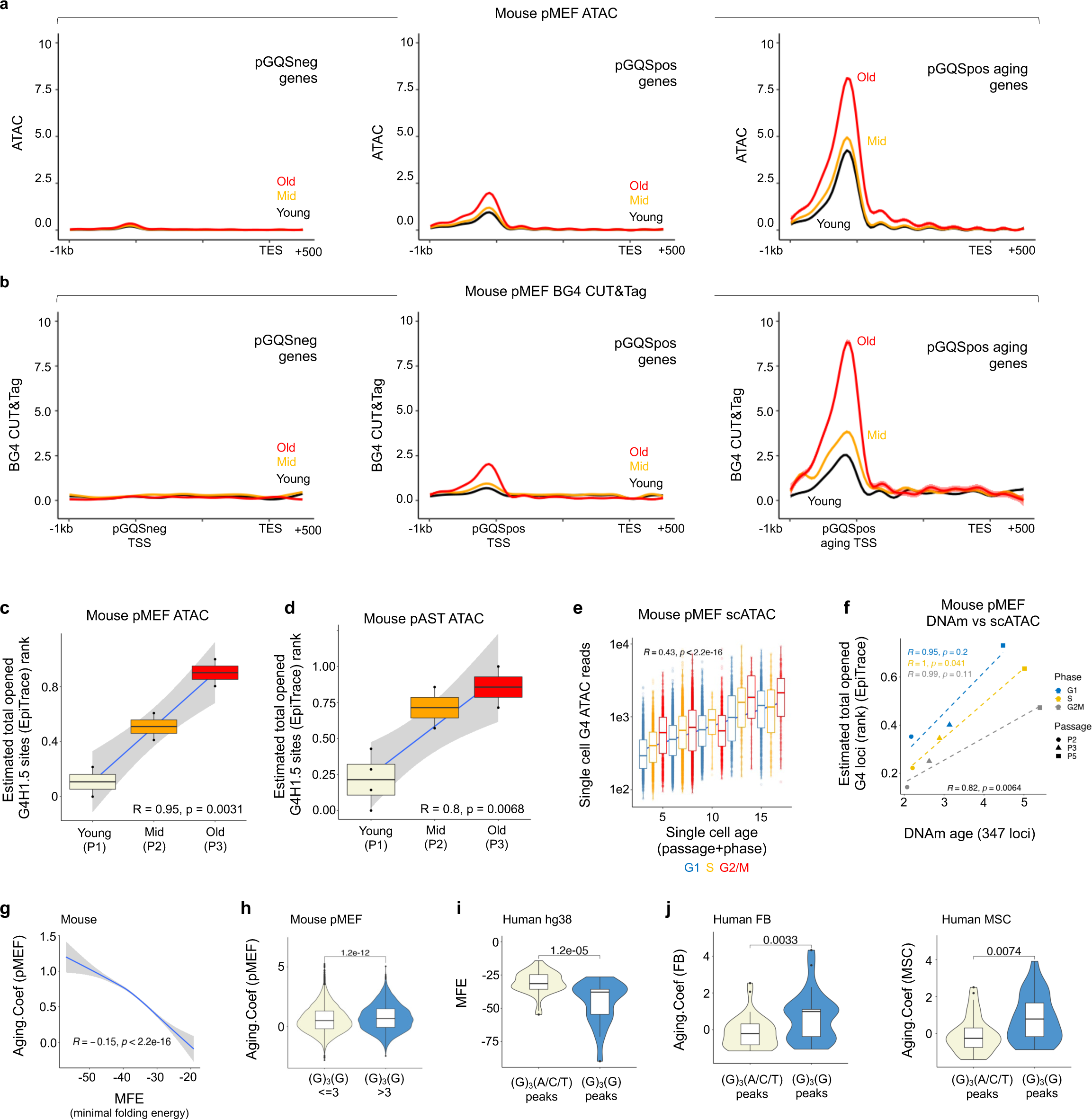
Clock-like formation of G4 associates with chromatin opening during aging. (a) ATAC-seq profile of young, mid-aged, and old pMEFs for genes without pGQS in their promoter (pGQSneg), genes with pGQS in their promoter (pGQSpos), and aging genes with pGQS in their promoter (pGQSpos aging). **(b)** BG4 CUT&Tag profile of young, mid-aged, or old pMEFs on the pGQSneg, pGQSpos, or pGQSpos aging genes. **(c)** Estimated total opened pGQS loci in bulk ATAC-seq of pMEFs. **(d)** Estimated total opened pGQS loci in bulk ATAC-seq of primary mouse astrocytes (pAST) cells. **(e)** Single-cell ATAC reads falling on the pGQS locus in pMEFs. Cells are ranked by their *in vitro* passage number and cell cycle phase (G1, S, and G2/M). **(f)** Correlation of average estimated total opened pGQS loci by EpiTrace from the scATAC data and the scAge-inferred DNA methylation age from the same batch of cells. Samples from the same cell cycle phases were subjected to linear regression. **(g)** Correlation between the G4 minimal folding energy (MFE) in the ATAC-seq peak (X-axis) and the aging coefficient of the same peak (Y-axis). **(h)** Aging coefficient of ATAC peaks containing <=3 or >3 G4 repeats. **(i)** MFE of human ATAC-seq peaks containing only G3/(A/C/T) repeats or containing only G3/(G) repeats. **(j)** Aging coefficients of human ATAC-seq peaks containing only G3/(A/C/T) repeats or containing only G3/(G) repeats in CiPSCs induced from fibroblasts (FBs) or mesenchymal stem cells (MSCs).

To test whether chromatin openings in G4 regions exhibit clock-like behavior, we used EpiTrace^7^ to estimate the total number of opened G4 loci in bulk or single-cell ATAC-seq data from *in vitro* passaged pMEFs or primary mouse astrocytes (pASTs). A linear correlation between the total number of opened G4 loci and cell age was found for different cell types across the different assays (Fig. 2c-e). Furthermore, the number of total opened G4 regions in the ATAC-derived sample correlated well with the DNAm-derived biological age of the pMEFs, regardless of the cell cycle phase (Fig. 2f).

The tendency of a particular locus to form a G4 is determined by the energy it releases during folding. We computationally inferred the minimal folding energy (MFE) of each putative G4 sequence by G4Boost^77,78^. In MEFs, the mean MFE of each aging locus correlated with the aging coefficient (Fig. 2g). In an orthogonal analysis, we classified the mouse aging loci by the number of stable (G3)/G repeats they harbored (Extended Data Fig. 2) and found that sites containing >3 (G3)/G repeats had a greater aging coefficient (Fig. 2h). Similarly, in the human genome, scATAC peaks containing (G3)/(G) repeats are thermodynamically more stable than are those containing (G3)/(A/C/T) repeats (Fig. 2i) and have a greater aging coefficient in two scATAC experiments (Fig. 2j-k). Hence, chromatin opening during aging is closely associated with G4 formation.

If G4 formation is necessary for clock-like chromatin opening, G4s should be enriched at aging loci compared to non-aging loci. To test this hypothesis, we computed loci-wise aging coefficients from mouse, nematode^79^, zebrafish^80^, Drosophila^81^ and human^22^ scATAC data and inferred the enriched DNA motives in the aging loci. The sequence motifs most strongly enriched in aging loci were generally G-rich sequences, many of which could form G4 structures (Fig. 3a, e, and Extended Data Fig. 3). In accordance with these findings, the pGQSpos genes had a much greater aging coefficient in their TSS than the pGQSneg genes did (Fig. 3a, e, and Extended Data Fig. 4). Furthermore, comparing to pGQSneg loci, the aging coefficient is increased in pGQSpos loci, and further increased in pGQS loci positive with G4 ChIP-seq signal (Fig. 3b, f) ^49,51,82^. Moreover, the pGQSneg locus harboring clock-like differentially methylated loci^8^ did not differ from the other pGQSneg loci (Fig. 3b, f). The presence of a G4 ChIP-seq signal correlated with the percentage of opened loci (Fig. 3c, g). These results indicate that G4 formation drives aging-associated, clock-like chromatin opening.

**Figure 3.**
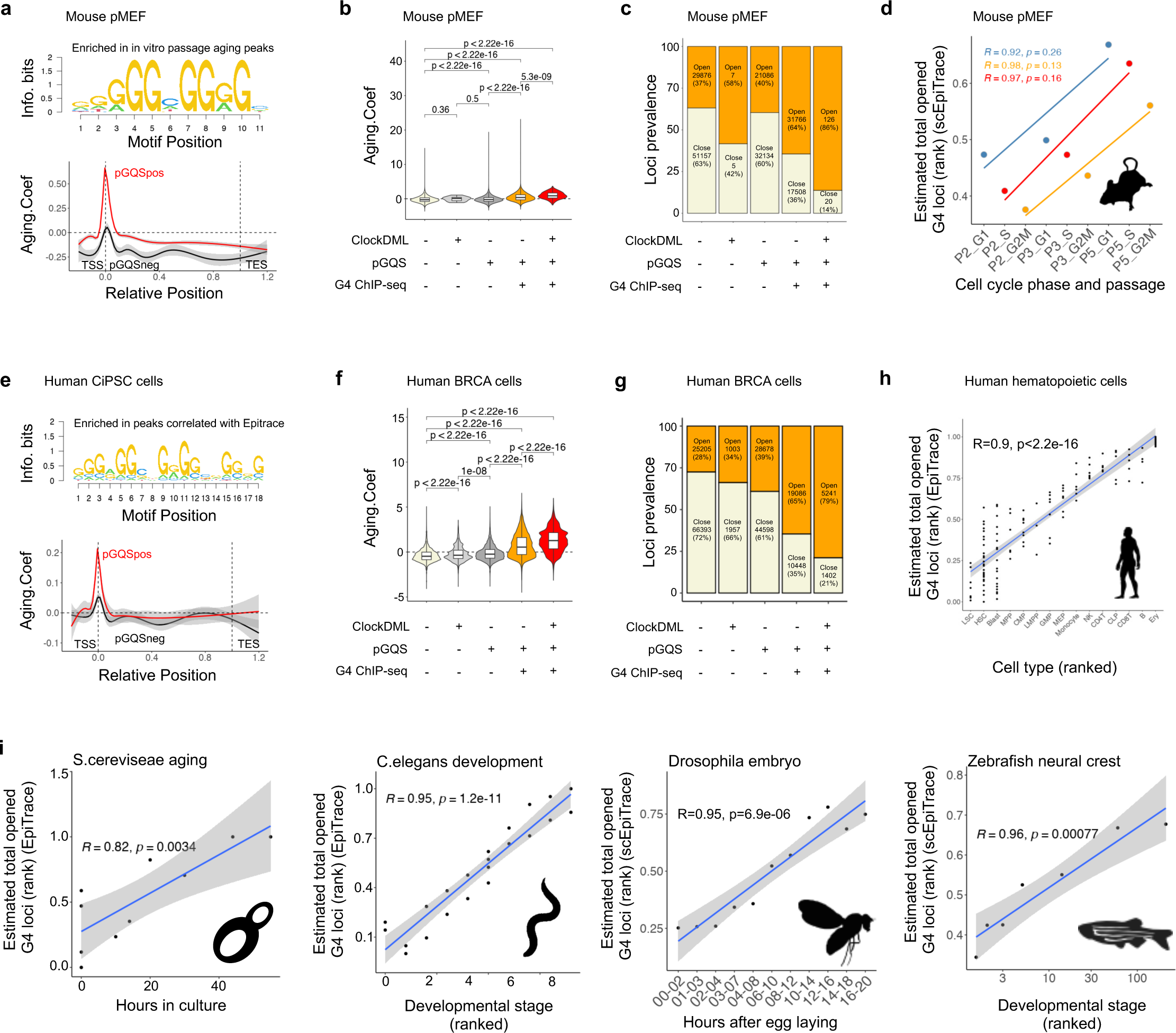
G4 formation is necessary for clock-like chromatin opening. (a) Top enriched sequence motif in aging ATAC-seq peaks in the mouse pMEF scATAC experiment (top) and aging coefficient profiles of the pGQSpos and pGQSneg genes in the same experiment (bottom). **(b)** Aging coefficients of ATAC-seq loci in mouse pMEF scATAC. Loci are classified by whether they harbor a ClockDML, a pGQS, or G4 ChIP-seq signal. **(c)** Relative prevalence of open (age >0, orange) and close (age <0, beige) ATAC-seq loci in mouse pMEF scATAC. Loci are classified by whether they harbor a ClockDML, a pGQS, or G4 ChIP-seq signal. **(d)** Correlation between the known sample age (X-axis) and the average estimated total number of opened G4 loci (Y-axis) in the mouse pMEF scATAC experiment. Samples from different cell cycle phases were separately analyzed. **(e)** Top enriched sequence motifs in aging ATAC-seq peaks in the human CiPSC scATAC experiment (top) and aging coefficient profiles of the pGQSpos and pGQSneg genes in the same experiment (bottom). **(f)** Aging coefficients of ATAC-seq loci in the human breast cancer PDX scATAC cohort. Loci are classified by whether they harbor a ClockDML, a pGQS, or G4 ChIP-seq signal. **(g)** Relative prevalence of open (age >0, orange) and close (age <0, beige) ATAC-seq loci in the human breast cancer PDX scATAC cohort. Loci are classified by whether they harbor a ClockDML, a pGQS, or G4 ChIP-seq signal. **(h)** Correlation of known cell developmental rank (X-axis) and the estimated total number of opened G4 loci (Y-axis) in the human hematopoietic cell ATAC experiment. **(i)** Correlations of known sample times (X-axis) and average estimated total number of opened G4 loci (Y-axis) from (1) the aging ATAC experiment, (2) the ATAC-seq data of *C. elegans* samples at known developmental stages, (3) the scATAC data of Drosophila samples at known developmental times, and (4) the scATAC data of zebrafish neural crest lineage cells at known developmental stages.

To test the universality of G4 locus chromatin opening during aging, we estimated the total number of opened G4 loci for accurately timed samples from mouse pMEF *in vitro* passages, from fly embryonic development^81^, from zebrafish neural crest development^80^, from *C. elegans* development^79^, from yeast experimental aging^83^, or from human hematopoietic cell development^84^ using EpiTrace. In all these cases, the estimated total number of opened G4 loci linearly correlated with known sample age (Fig. 3d, i) or known rank of cell development (Fig. 3h) with high accuracy. These data indicate that gradual opening of G4 loci is a common feature of aging throughout evolution.

### Pioneer factor primes G4 formation on nucleosome-depleted regions of promoters

To understand the molecular mechanism underlying the aging-associated accumulation of G4s and associated chromatin opening, we first searched for potential molecular initiators of G4 formation via bioinformatics. We performed unbiased statistical enrichment for published TF ChIP-seq data in pGQS^57^ in the mouse and human genomes^85^. As reported, many TFs are strongly enriched in pGQS^86^. Interestingly, the most strongly pGQS-enriched TFs in mice (Extended Data Fig. 5a) or humans (Extended Data Fig. 5b) are pioneer factors that bind to tightly packed, nucleosome-wrapped DNA and open chromatin^35^. In mice, three out of four “Yamanaka factors” for pluripotency induction^21^ and two auxiliary pluripotency TFs (Ell3^87^ and Prdm14^88^) composed 71% (5/7) of the selected top pGQS-enriched (top 10% q-value) TFs (Extended Data Fig. 5a). In humans, the known pioneer factors FOXA2, GATA2 and SPI1/PU.1 and ZBTB7A, the homolog of the *Drosophila* pioneer factor GAF^89^, are among the top 10 pGQS-enriched TFs (Extended Data Fig. 5b).

We tested the interdependence of pioneer factor binding and G4 formation on pGQS by analyzing ATAC-seq and RNA-seq data from mouse cells with genetic perturbation of the pioneer factors Oct4, Nanog, Sox2^90^, and Zbtb7a^91^. ChIP-seq of these TFs revealed the expected enrichment of the pGQSpos gene TSS compared to the pGQSneg gene TSS (Extended Data Fig. 6-8). On the other hand, BG4 CUT&Tag signals are enriched on TSSs with pioneering factor binding sites (Extended Data Fig. 8). Although knocking down Oct4, Sox2, or Nanog does not affect chromatin accessibility at G4 loci (Extended Data Fig. 6), Zbtb7a knockout results in a reduction in chromatin accessibility at pGQSpos (Extended Data Fig. 8). Similarly, Zbtb7a knockout affects the opening of the pGQSpos and pGQSneg TSSs and decreases the 5’ RNA expression of their bound targets, regardless of whether they harbor a pGQS in the TSS (Extended Data Fig. 8), indicating decreased occupation of paused RNA polymerase II at these sites. Hence, the pioneer factor Zbtb7a binds to pGQS prior to G4 formation to create a nucleosome-depleted region (NDR), which enables G4 formation.

### G4 stimulates transcription-replication interactions to delay genome replication

The pGQSpos TSS, particularly the pGQSpos TSS of aging genes, is enriched with RNA polymerase II Ser5 phosphorylation and NELF, two essential drivers of transcriptional elongation^92–94^ (Fig. 4a and Extended Data Fig. 9). DNA replication origin activity^95^ was also enriched on the pGQSpos TSS compared to the pGQSneg TSS (Fig. 4e-f). The application of G4-binding small molecule, which potentially impairs the recruitment of proteins to G4^93,94^, negatively affects both transcriptional elongation, as shown by nascent RNA sequencing (Fig. 4b), and DNA replication on the pGQSpos TSS (Fig. 4e), indicating that G4 formation stimulates both transcriptional elongation and DNA replication in its vicinity.

**Figure 4.**
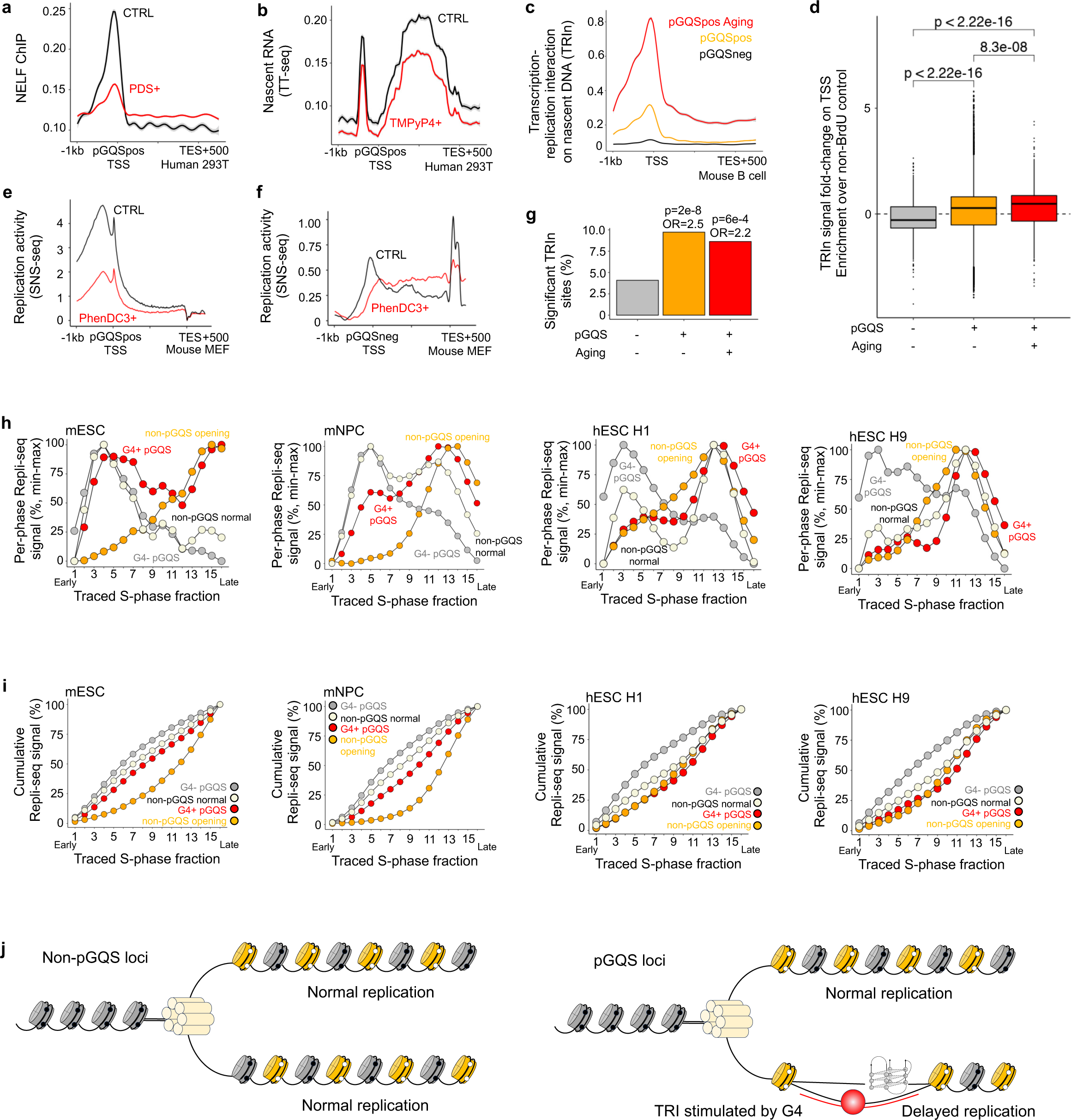
G4 promotes transcription-replication interactions to delay genome replication at aging loci. (a) NELF ChIP-seq profiles of control (CTRL) and PDS-treated (PDS+) HEK293T cell samples for pGQSpos genes. **(b)** Nascent RNA (TT-seq) profiles of control (CTRL) and TMPyP4-treated (TMPyP4+) HEK293T cell samples of the pGQSpos gene. **(c)** Profiles of transcription-replication interactions on nascent DNA (TRIPn) on the pGQSneg, pGQSpos, and pGQSpos aging genes in mouse spleen B cells. **(d)** Sitewise TRIPn signal enrichment (over non-BrdU control) of the pGQSneg, pGQSpos, and pGQSpos aging genes in mouse spleen B cells. **(e)** Short replication-associated nucleotide (SNS-seq) profiles of control (CTRL) and PhenDC3-treated (PhenDC3+) mouse MEF cell samples of the pGQSpos gene. **(f)** Short replication-associated nucleotide (SNS-seq) profiles of control (CTRL) and PhenDC3-treated (PhenDC3+) mouse MEF cell samples of the pGQSneg gene. **(g)** Percentage of mouse spleen B cells harboring significant enrichment of the TRIPn signal from the pGQSneg, pGQSpos, and pGQSpos aging genes. **(h)** BrdU pull-down Repli-seq signal across 16 fractions (time points) of S phase cells grouped according to different classes of genes in mouse embryonic stem cells (mESCs), mouse neural progenitor cells (mNPCs), and human embryonic stem cells (hESCs, H1 and H9). **(i)** Cumulative BrdU pull-down Repli-seq signal (normalized to min(0%)-max(100%)) across 16 fractions (time points) of S phase, grouped by different classes of genes, in mouse embryonic stem cells (mESCs), mouse neural progenitor cells (mNPCs), and human embryonic stem cells (hESCs, H1 and H9). **(j)** Schematic of the finding. G4 stimulates transcription-replication interactions by simultaneously promoting both transcription and replication, resulting in delayed genome replication.

Strong transcriptional activity coupled with DNA replication activity could enhance transcription-replication interaction (TRI), stimulating the formation of DNA-RNA hybrids that might hinder DNA replication progression^59,63,64,96^. We found that elongating RNA polymerase II-bound newly synthesized DNA^97^ is significantly enriched on the pGQSpos TSS (Fig. 4c-d). This enrichment is even more pronounced at the TSS of aging genes associated with pGQSpos (Fig. 4c-d). As a result, significantly more transcription-replication interaction sites were found for the pGQSpos genes (Fig. 4g). In accordance with these results, aging coefficients across the genome are associated with the TRI tendency^98^ (Extended Data Fig. 10). These data indicate that G4 formation stimulates TRI by simultaneously promoting both transcription elongation and DNA replication.

The transcription-replication interaction presents a roadblock for DNA replication, which must be resolved before replication resumes. G4-mediated blockade of DNA replication reportedly delays DNA replication in chicken REV-1-deficient cells^99^. To test whether G4-mediated blockade of DNA replication is universal, we analyzed genome replication dynamics in four different mouse and human cell lines with high-resolution Repli-seq^100^ data (Fig. 4h-i). In all the cell lines, comparison of the G4 ChIP-seq-negative pGQS loci to the G4 ChIP-seq-positive pGQS loci revealed that the presence of G4 strongly delayed DNA synthesis (Fig. 4h-i). In non-pGQS loci, age-associated chromatin opening is also associated with delayed DNA synthesis (Fig. 4h-i). Taken together, these data indicate that delayed genome replication is a general feature of aging loci and that G4 stimulates local TRI to delay genome replication (Fig. 4j).

### G4s impair DNA methylation transmission during aging

Delayed genome replication shortens the available time for histone modification recovery and hinders parental histone recycling to the newly synthesized daughter chromosome. Therefore, we hypothesize that impaired epigenomic transmission from the parental to daughter genome results from delayed genome replication at G4 loci.

We first tested whether the transmission of DNA methylation from the parental to daughter genome is affected by G4. To analyze DNA methylation transmission, we analyzed DNA methylation maintenance between parental and daughter DNA strands in human HeLa cell Hammer-seq data^101^.

As a positive control of the analysis, DNA methylation maintenance was impaired at the solo-WCGW site compared to the non-WCGW and non-pGQS control CpG sites^101^ (Fig. 5a). Although the non-G4 pGQS sites did not differ from the control sites, the G4 sites exhibited significantly delayed DNA methylation maintenance in both the early and late S phases (Fig. 5a). This phenomenon was validated by analyzing another Repli-BS-seq dataset^102^ (Extended Data Fig. 11). The difference in DNA methylation maintenance efficiency between the G4 locus and the non-G4 pGQS locus diminished upon double knockout of DNMT3A/3B, indicating that G4 formation hinders DNMT3A/3B-mediated DNA methylation (Extended Data Fig. 11).

**Figure 5.**
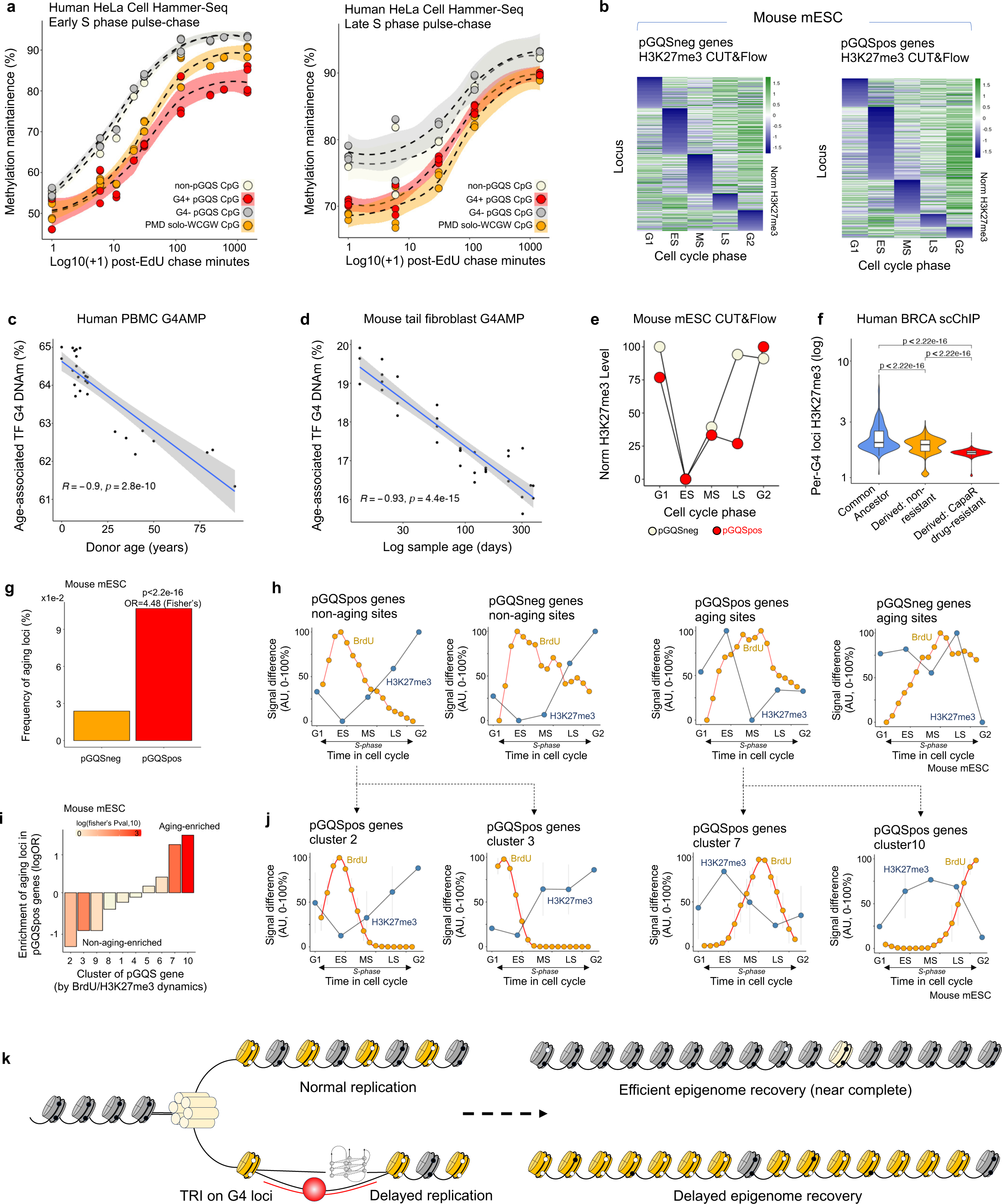
Delayed genome replication by G4 impairs epigenome transmission during aging. (a) Methylation maintenance level of early (left) or late (right) S-phase EdU pulse-thymidine-chase HeLa cells, nonpGQS CpG, PMD solo-WCGW CpG, G4-pGQS CpG, or G4+ pGQS CpG according to Hammer-seq experiments. The post-EdU chase times are shown on the X-axis (log-ed). **(b)** Normalized H3K27me3 CUT&Flow signals of pGQSpos (left) and pGQSneg (right) in mouse mESCs. **(c)** DNA methylation level at age-associated TF-bound G4 peaks in human PBMC G4AMP samples. Human donor ages are shown on the X-axis. **(d)** DNA methylation level at age-associated TF-bound G4 peaks in mouse tail fibroblast G4AMP samples. Mouse sample ages are shown on the X-axis (log-ed). **(e)** Mean H3K27me3 CUT&Flow signal profiles of pGQSpos (left) and pGQSneg (right) mouse mESCs across different cell cycle phases. **(f)** Single-cell H3K27me3 ChIP-seq signal per G4 locus in human breast cancer PDX cells. The cells were clustered into common ancestor cells, derived cells in non-drug-resistant sample, and derived cells in the drug-resistant sample. **(g)** Relative frequency (%) of pGQSneg or pGQSpos ATAC peaks exhibiting an aging peak. **(h)** Average DNA replication (BrdU, yellow) and normalized H3K27me3 signal (H3K27me3, blue) across different time points (X-axis) during cell cycle progression. **(i)** Relative enrichment of aging-related loci in the pGQSpos gene cluster. These pGQSpos genes are clustered according to their Repli-seq and H3K27me3 CUT&Flow temporal profiles. **(j)** Average DNA replication (BrdU, yellow) and normalized H3K27me3 signal (H3K27me3, blue) across different time points (X-axis) for non-aging and aging pGQSpos genes. **(k)** Schematic of the findings. During DNA replication, G4 might directly hinder the replication fork or create TRI in nascent synthesized DNA, resulting in delayed genome replication. Such delayed genome replication reduced the available time window for histone modification and/or G4-associated DNA methylation recovery.

To validate whether DNA methylation transmission is impaired by G4 formation during aging, we analyzed DNA methylation levels around G4 loci in mouse and human cells by targeted bisulfite sequencing with anchored PCR (G4AMP or MArchPCR). In mouse and human primary cells, CpG in the vicinity of G4 could undergo hypomethylation during aging (Extended Data Fig. 12a, b, e, h).

Since not all CpGs in the vicinity of G4 are similarly hypomethylated during aging and since the thermostability of G4 is correlated with age-associated chromatin opening (Fig. 2g-k), we hypothesized that G4 stability might also regulate age-associated DNA hypomethylation. Since G4 is a hub for TF binding^55,86^, we tested whether TF activity could regulate G4-associated DNA hypomethylation. We inferred TF activity during pMEF *in vitro* passage using SHARE-seq data^103,104^ and determined three TFs (Zfp148, Zfp263, and Klf15) that significantly associated with age (Extended Data Fig. 12c). We then analyzed the DNA methylation level of the same sample CpGs in the vicinity of G4 bound by these TFs. Interestingly, the CpGs falling in these regions showed consistent age-associated DNA hypomethylation (Extended Data Fig. 12d). Hence, G4 DNA hypomethylation during aging is regulated by TF activity.

We then classified G4 regions according to the presence of TF-binding sites (TFBS) and separately analyzed DNA methylation in each group. In both mouse and human cells, most G4-bound TFs are associated with DNA hypomethylation during aging (Extended Data Fig. 12f, g, i, j). Overall, in the vicinity of G4 bound by these TFs, aging was associated with a strong and linear decrease in the DNA methylation level (Fig. 5c-d). Taken together, these results indicate that G4 impairs DNA methylation transmission during aging.

### G4 impairs histone modification transmission during aging

To analyze the effect of G4 on histone modification transmission, we first analyzed histone modification recovery using mouse ESC H3K27me3 CUT-and-flow data^105^ (personal communication). We calculated the frequency of gene-associated H3K27me3 modifications by averaging H3K27me3 CUT&Tag signals from the flanking +/-50 kb regions around the gene. At each locus, the H3K27me3 signal valley across the cell cycle indicates the completion of histone exchange, and its peak indicates the completion of recovery of the parental H3K27me3 modification. In general, the pGQSpos gene is more likely to undergo complete histone exchange in the early S phase (Fig. 5b and Extended Data Fig. 13a) and requires slightly more time to recover H3K27me3 modification (Extended Data Fig. 13b-c). As a result, the mean H3K27me3 recovery in the pGQSpos genes was slower than that in the pGQSneg genes (Fig. 5e).

We validated whether H3K27me3 at G4 loci is reduced during cell aging using single-cell H3K27me3 ChIP-seq (scChIP) data derived from a human breast cancer (BRCA) PDX model^106^. In this study, BRCA cells (HBx95) were grafted into mice and subjected to chemotherapy (capecitabine). Clustering of the scChIP profiles of cancer cells from different samples revealed a common ancestor cell population that existed in both the drug-treated and control samples (Extended Data Fig. 14a-c) and two derived cell populations corresponding to either the control, non-drug-resistant, or drug-resistant state (Extended Data Fig. 14c). Compared to those of the ancestor population, the H3K27me3 coverage of the G4 locus decreased (Fig. 5f) regardless of the drug resistance status, and the H3K27me3 coverage of the G4 locus further decreased (Fig. 5f and Extended Data Fig. 14d). These results indicate that G4 impairs histone modification transmission in its vicinity.

### Aging-associated chromatin opening results from impaired epigenome transmission

Impaired epigenome transmission by G4s during each cell cycle results in gradual DNA hypomethylation and loss of heterochromatin (Fig. 5a-f). This in turn would create a more permissive local environment for G4 formation in subsequent generations. This self-reinforcing cycle of G4 formation not only explains the gradual accumulation of G4s during aging (Fig. 1) but also suggests concurrent clock-like epigenomic changes in the neighboring G4 genomic region.

Aging peaks were more enriched at the pGQSpos locus than at the pGQSneg locus (Fig. 5g). Analysis of DNA replication and histone modification recovery dynamics revealed that in non-aging sites, DNA replication is completed in the early S-phase, allowing long recovery times for histone modification (H3K27me3; Fig. 5h). This finding is in stark contrast to what has been observed in aging sites, where DNA replication does not complete until late S-phase, and histone modification recovery is delayed until the next cell cycle (Fig. 5h). We then clustered the pGQSpos genes based on their DNA replication and H3K27me3 recovery dynamics using K-means clustering on time series^107^ (Fig. 5i) and found a similar scenario: aging sites were depleted from the early replicating clusters and enriched in the late replicating clusters that could not finish H3K27me3 recovery within the cell cycle (Fig. 5j). Taken together, these results indicate that G4 impairs epigenome transmission across mitosis to gradually erode the local epigenetic landscape (Fig. 5k).

### Genetic mutations causing G4 accumulation accelerate the erosion of the epigenetic landscape around G4 *in vivo*

Mutations in G4-resolving enzymes such as BLM, WRN, ERCC6 and the E3 ligase ERCC8 result in premature aging in human patients. Previously, it was reported that mutations in WRN result in global loss of H3K9me3 in human mesenchymal stem cells (hMSCs) ^108^, but there is no clear link between biochemical function and phenotype. We reanalyzed the H3K9me3, H3K4me3 and H3K27me3 ChIP-seq data from this study. Strikingly, the WRN mutation resulted in a much greater reduction in H3K9me3 at putative G4 loci than at non-pGQS loci (Fig. 6a). Although the WRN mutation induces only a minor and local reduction in H3K9me3 at non-pGQS loci, it induces a potent reduction in H3K9me3 at pGQS loci (Fig. 6a). This effect is not limited to the local vicinity of pGQS and can extend to distal sites as far as 50 kb away (Fig. 6a). Furthermore, the WRN mutation induces a gain of H3K4me3 specifically at the pGQS locus (Fig. 6b) and decreases H3K27me3 peaks globally (Fig. 6c).

**Figure 6.**
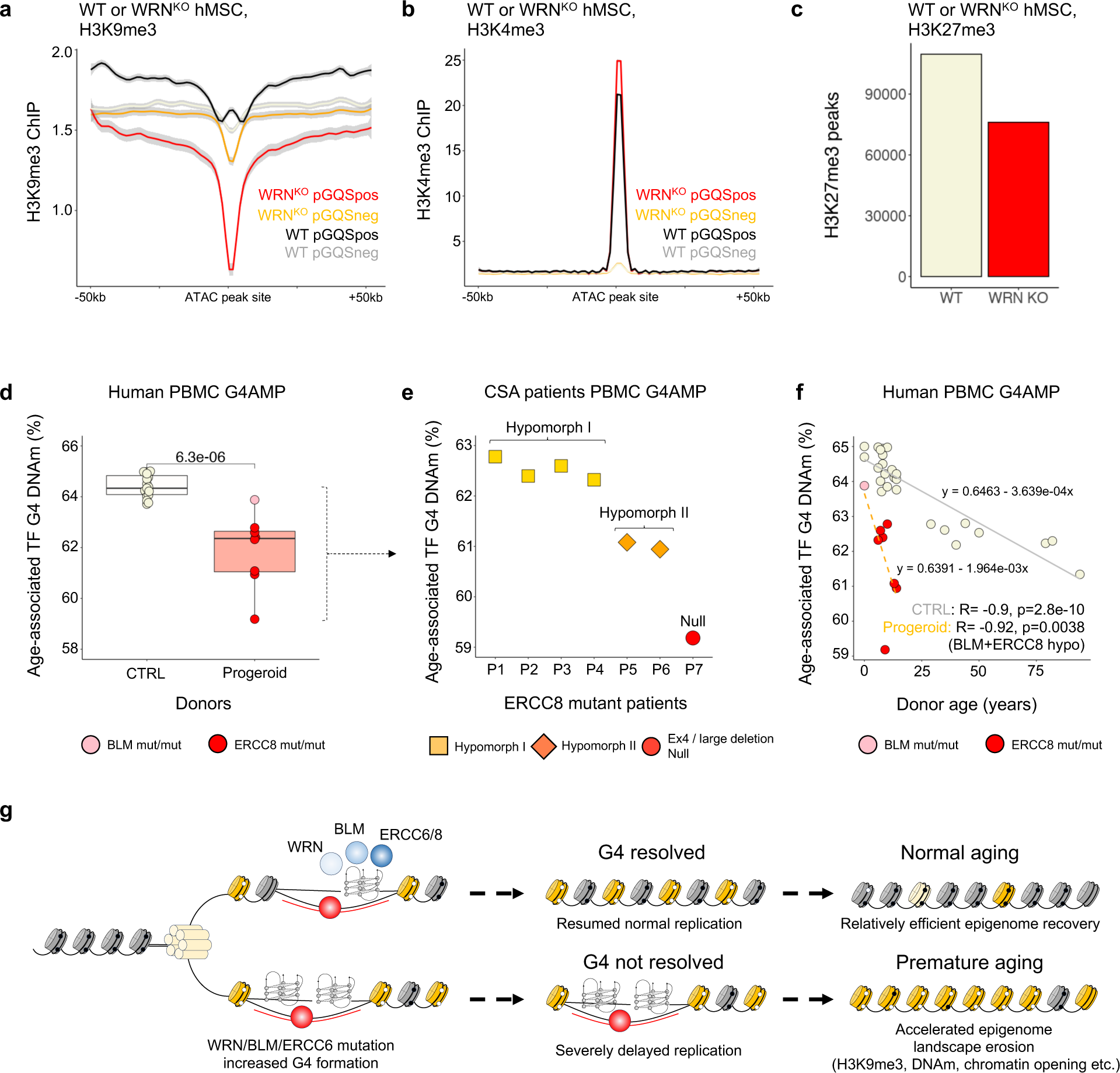
Genetic mutations causing G4 accumulation accelerate the erosion of the epigenetic landscape around G4 *in vivo*. (a) H3K9me3 profiles across the pGQSpos or pGQSneg sites. Profiles were determined between +/− 50 kb of a particular given site for wild-type (WT) or WRN knockout (WRN KO) hMESC samples. **(b)** H3K4me3 profiles across the pGQSpos or pGQSneg sites. Profiles were determined between +/− 50 kb of a particular gene in wild-type (WT) or WRN knockout (WRN KO) hMESC samples. **(c)** Number of H3K27me3 peaks from wild-type (WT) or WRN knockout (WRN KO) ATAC samples. **(d)** DNA methylation level at age-associated TF-bound G4s peaked in age-matched progeroid and normal control donors. **(e)** DNA methylation level at age-associated TF-bound G4 peaks in ERCC8-mutant progeroid patients segregated by the nature of the ERCC8 mutation (hypomorphic or null). **(f)** Linear regression of DNA methylation levels in age-associated TF-bound G4 peaks in ERCC8-mutant patients and controls (Y-axis) according to donor age (years, X-axis). **(g)** Summary of findings. WRN/BLM/ERCC8 mutations result in accelerated erosion of the epigenetic landscape adjacent to G4.

We analyzed G4-associated DNA methylation by sequencing PBMCs from a group of human progeroid patients with mutations in BLM or ERCC8. Significant aging-associated G4 DNA hypomethylation was detected in the progeroid patients compared to the age-matched non-progeroid donors (Fig. 6d). Furthermore, segregating the ERCC8 patients into groups according to the molecular mutations they carried (Extended Data Fig. 15a) revealed that the aging-associated G4 DNA hypomethylation level was correlated with mutation severity: the DNA hypomethylation phenotype was least severe in patients with compound heterozygous ERCC8 mutations with a hypomorphic short nucleotide variation mutation over a structural variation^109^ and most severe in a patient with structural variation over a genetically null large chromosomal deletion (Fig. 6e). Compared to that in non-progeroid donors, DNA hypomethylation around G4 in these patients accelerated to such an extent that a 10-year-old patient had a DNA methylation level way below that of a 94-year-old non-progeroid donor (Fig. 6f). Linear regression shows that in these patients, the aging-associated G4 DNA hypomethylation was accelerated by a factor of 5 compared to controls, which agrees well with the usual life expectancy of 10∼20 years of Cockayne and Bloom syndrome patients (Fig. 6f). Analysis of the full pGQS DNA methylation level revealed similar findings (Extended Data Fig. 15b-d). To summarize, these data indicate that pathogenic mutations causing G4 accumulation accelerate the progression of the epigenetic landscape around G4 *in vivo*.

## Discussion

Aging is a fundamental and universal phenomenon of life. Here, we showed that G4 formation drives aging in eukaryotes by promoting the erosion of the epigenetic landscape (Fig. 7). Pioneer factor primes G4 formation on nucleosome-depleted regions of the transcription start site. Once G4s are formed, they stimulate transcription-replication interactions, which delay genome replication. Delayed genome replication results in impaired epigenome transmission. As a result, G4 triggers the gradual erosion of its neighboring epigenetic landscape.

**Figure 7.**
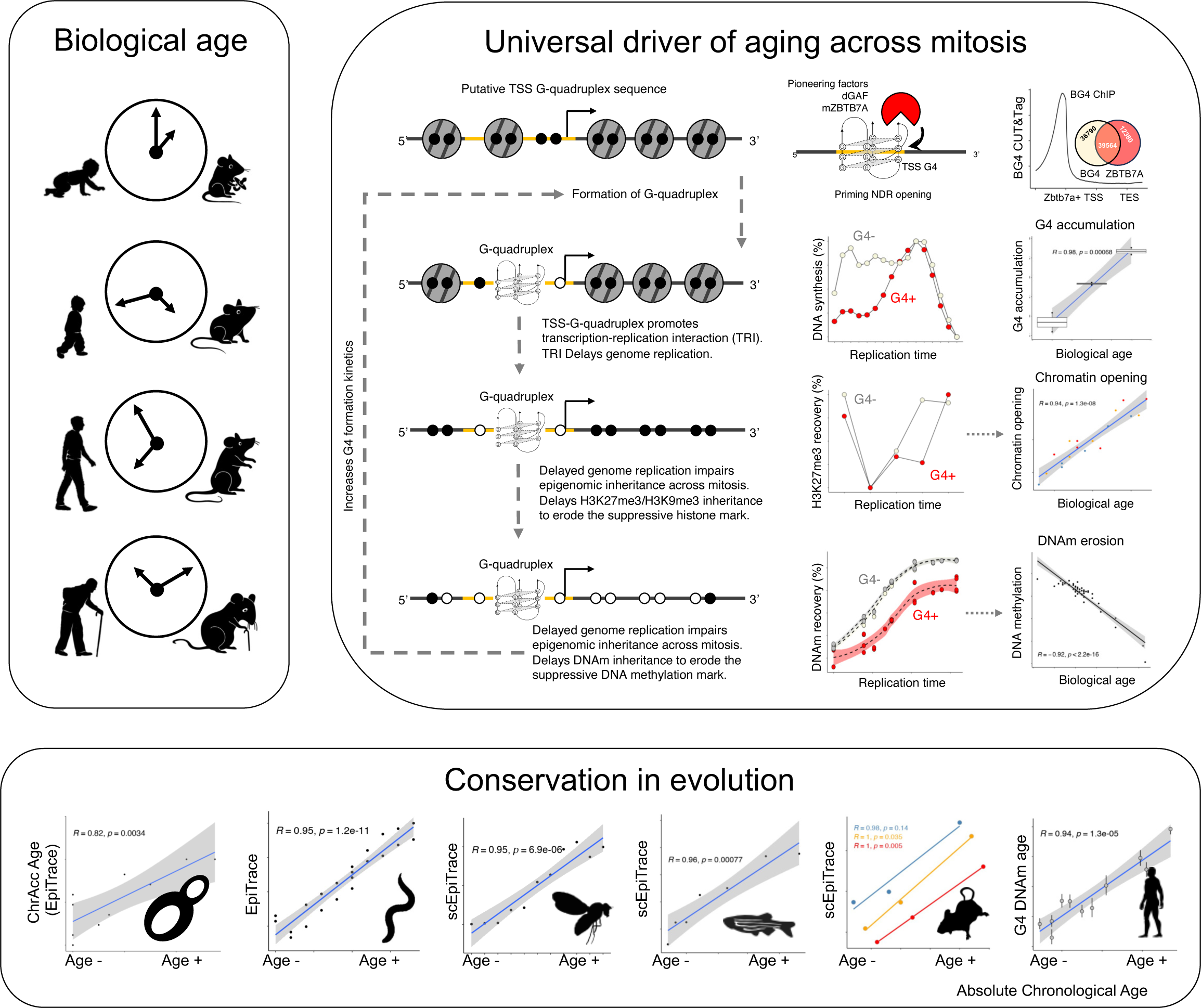
Summary of G4-driven aging. During eukaryotic cell aging, putative G-quadruplex regions at transcription start sites can be opened by pioneering factors such as Zbtb7a/GAF. The opened chromatin stochastically folds into a G-quadruplex. The G-quadruplex induces transcription-replication interactions to delay genome replication in S phase and impairs epigenetic information transmission locally, resulting in gradual, clock-like DNA hypomethylation and chromatin opening. G-quadruplex-driven epigenomic aging is highly conserved during evolution.

It is natural to presume that G4s form in open chromatin spontaneously. However, the accumulation of G4s across mitosis is not uniform across the genome and might differ between cell types. Through bioinformatic analysis, we determined that this accumulation is associated with G4 thermostability and might be modulated by protein binding. Identifying the exact molecular mechanism of aging-associated, cell type-specific G4 accumulation is of particular interest for understanding aging-related diseases affecting specific cell types, such as arteriosclerosis, Alzheimer’s disease, and amyotrophic lateral sclerosis.

Immunostaining of DNA G4s using a single-chain antibody revealed spot-like G4s in the nucleus^48^. However, G4-associated epigenetic changes, including H3K9me3 erosion and DNA hypomethylation in heterochromatin domains, suggest that these events initially occur near the nuclear lamina. Human mutations in nuclear lamina proteins result in premature aging and inactivation of normally silenced LINE1 retrotransposons in heterochromatin domains^110^. In *Drosophila*, GAF, the *Drosophila* homolog of Zbtb7a^89^, binds to a GAGA-rich sequence that is enriched according to ChIP-seq data of the *Drosophila* nuclear pore basket protein Elys^111^.

Interestingly, our results also revealed the GAGA-rich sequence as the most strongly enriched motif at loci involved in aging-associated chromatin opening in the *Drosophila* genome. Furthermore, the pathogenic RNA G4 structure in C9orf72 causes neurodegeneration through blocking nuclear pore trafficking^112,113^, indicating an intimate link between the nucleic acid G4 structure and the periphery of the nucleus. Although the exact physical location of initial G4 formation during aging is currently unknown, we consider this a potentially interesting research direction.

G4s exist in multiple forms. In particular, pathogenic mutations in progeroid disease might affect different types of G4s: while WRN and BLM mostly act on intrastrand G4s^44^, ERCC6 specifically binds to interstrand G4s and might mediate their resolution^43^. Interestingly, WRN-knockout hMSCs exhibit minimal differences in DNA methylation compared with wild-type hMSCs^108^, while large shifts in H3K9me3 and H3K4me3 occur around G4 loci. On the other hand, our results showed that compared with those from controls, PBMCs from ERCC8-mutated patients exhibit prominent G4 DNA hypomethylation. It would be interesting to test whether different types of G4s cause different types of epigenetic inheritance deficiency.

In the mammalian system, inducing constitutive faithful DNA repair alone causes global epigenomic information attrition and organismal aging^25^. G4 can promote replication-dependent genome instability^65^ and is enriched in DNA breakpoints in cancer^58^. Since all progeroid-pathogenic G4-resolving helicases function in DNA repair, G4 accumulation may promote aging via a DNA-break-repair mechanism. Although our study does not exclude this possibility, the localized effect of G4 on H3K9me3 and DNA methylation loss in WRN and ERCC8 mutants suggested that replication-dependent epigenomic transmission error is a more likely explanation for these results.

Nonetheless, our study is not without limitations. The application of G4-binding small molecules might affect both the stability of G4 and the binding of G4 to its interaction partners^93,94^; hence, G4 could be either antagonistic or agonistic in nature. Therefore, the results derived from G4-binding small molecule manipulations could have alternative explanations. Since there are no straightforward methods for affecting G4 stability without simultaneously perturbing its binding with partners, such limitations seem unavoidable. Finally, our work focused on the G4 formation cycle throughout mitosis. However, aging also occurs in cells that do not actively divide, such as neurons. The chromatin accessibility of neurons shifts during aging^114^, suggesting that erosion of the epigenetic landscape might also underlie aging in these cells. However, whether G4s might actively regulate aging in these non-dividing cells, and if so, what the underlying molecular mechanism is, is an open question awaiting future investigation.

In conclusion, our results revealed a highly conserved molecular mechanism driving eukaryotic aging. G4-resolving enzymes were previously proposed to be “mitotic counters” of a cell^69^ since cells harboring their mutations have a shortened Hayflick limit. Through our analysis, we established a molecular link to explain how pathogenic mutations in G4-resolving enzymes accelerate mitosis to cause human progeroid disease. While analyzing G4-associated epigenetic changes could serve as a measure of biological age, perturbing G4 formation might be of particular interest for modulating natural and pathological aging.

## Methods

### Human donors, animal samples and cells

This study was conducted in accordance with local legislation, measures of the Declaration of Helsinki, and the Ethics Protocols of the Human Genetic Resource Preservation Center of Hubei Province, China (Hubei Biobank). The human samples used in this study were pretreated and preserved by the Hubei Biobank, approved by the Institutional Ethical Review Board (approval number: 2017038-1). Informed consent was obtained from all individuals. For the mouse experiments, 33 C57BL/6 mice were used for tail fibroblast collection, 5 pregnant C57BL/6 mice were used for primary embryonic fibroblast collection, and 3 dams of C57BL/6 neonatal mice were used for primary astrocyte collection. The animal experiments were carried out at Zhongnan Hospital of Wuhan University and were approved by the Institutional Ethical Review Board (approval number: ZN2023148).

### Datasets used in this study

A full list of public datasets used in this study is shown in Extended Data Table 1.

### Genotyping of progeroid patients

The patients were initially subjected to whole-exome sequencing, and the candidate pathogenic mutations were confirmed by Sanger sequencing. The ERCC8 patients were further genotyped as described in Wang *et al*., 2017^115^. The ERCC8 exon 4 complex rearrangement was sequenced as described by Ren *et al*., 2003^116^. A full list of the genotypes of these patients is shown in Extended Data Table 2.

### Cell culture

Human bone marrow-derived mesenchymal stem cells (BMSCs) were obtained from the Stem Cell Bank, Chinese Academy of Sciences. Human BMSCs were cultured in MSC NutriStem® XF Medium (Biological Industries, 05-200-1A) supplemented with MSC NutriStem® XF Supplement Mix (Biological Industries, 05-201-1U) and 1% penicillin and streptomycin. The cell handling procedures were conducted according to the instruction manual. Primary mouse embryonic fibroblasts (MEFs) were isolated and maintained following a previously described protocol^117^. The FACS-sorting and SHARE-seq sequencing methods used for pMEFs were described in a previous study^7^.

### BG4 antibody expression and purification

pSANG10-3F-BG4^48^ (Addgene #55756) was subsequently transformed into BL21 cells. A single clone of the transformant was precultured in 3 mL of 2xTY + 2% glucose medium at 30°C overnight and subsequently transferred to 1 L of ZYM-5052 autoinduction medium^118^ for culture at 25°C overnight. Bacteria were collected by centrifugation. Osmotic extraction was performed by resuspending the bacteria in 20 mL of TES^119^ with benzonase (Millipore #E1014) on ice for 1 hr, followed by centrifugation at 10,000xg for 10 mins at 4°C. Supernatants were collected and diluted 1:10 into PBS with 10 mM NaCl, pH 8.0, passed through 0.22 µm filter, captured on column with 1 mL cobalt resin (Takara #635504), washed by 15 column volumes of PBS with 100 mM imidazole and 100 mM NaCl, pH 8.0, then eluted by 3 column volumes of PBS with 200 mM imidazole and 100 mM NaCl, pH 8.0. The collected proteins were concentrated by an Amicon Ultra-3k centrifugal filter (Millipore), exchanged with PBS supplemented with 100 mM NaCl and 50% glycerol, and stored at −80°C.

### BG4 immunostaining and automatic quantification

BG4 immunostaining was performed as previously described^48^. Briefly, live MEFs at different passages were grown on glass coverslips, fixed in methanol:acetic acid (3:1) and permeabilized with 0.1% Triton-X100/PBS. After blocking in 2% BSA/PBS, immunofluorescence was performed using standard methods with BG4 (1:50), anti-FLAG (Sigma, F1804, 1:100) and Goat anti-Mouse, Alexa Fluor 594 (Invitrogen, A-11032, 1:1000) antibodies. Coverslips were mounted with Prolong Gold/DAPI (Invitrogen). Quantification was performed in R (4.1.3) using the package EBImage (4.42.0) programatically, without human intervention, to select for ROIs or adjust intensities.

### Chromatin accessibility profiling (ATAC-seq) and CUT&Tag sequencing

CUT&Tag and ATAC-seq experiments were performed according to the protocol used in Xiao *et al*., 2022^120^. For the quantitative ATAC-seq experiments, 20,000 human HEK293T cells (kindly provided by Dr. Qi Li, National Institute of Biological Sciences, Beijing, China) were mixed with 50,000 mouse cells prior to the experiment. For quantitative CUT&Tag experiments, 40,000 human HEK293T cells were mixed with 70,000 mouse cells prior to the experiment. The antibodies used in the CUT&Tag experiments were BG4 (homemade), mouse anti-FLAG (Sigma, # F1804), and goat anti-mouse IgG (Sangon, #D111024-0001). Fragmentation after antibody binding was performed with a NovoProtein CUT&Tag 2.0 pAG-Tn5 kit (NovoProtein Ltd., Cat. #N259).

### qPCR of G4 enrichment in the opened region

CUT&Tag NGS libraries were amplified with (1) a G-quadruplex-specific common primer (NGS-1290, 5’-<u>GGAGGGGAGGGGAGAGGGG</u>-3’) and the Illumina P7 adaptor common primer or (2) a pair of Illumina P5/P7 common primers as controls for DNA content in qPCR mix (VAHTS SYBR qPCR master mix (Vazyme, #NQ101) on a Quantagene q225 qPCR machine (Kubo Technologies, Beijing, China). Amplification curves were analyzed by the standard ddCt method.

### Anchored PCR bisulfite sequencing

The multiplex anchored PCR bisulfite sequencing data of mouse primary embryonic fibroblasts generated using the MM285/347 ClockDML primer set were obtained from a previous publication^7^. For G4-specific DNA methylation sequencing (G4AMP) of human or mouse samples, genomic DNA was extracted from the proteinase K-digested tail tips of mouse or peripheral blood mononuclear cell (PBMC) of human donors using DNeasy blood tissue kit (Qiagen, #69506). 25 ng of genomic DNA was subjected to bisulfite conversion using a commercial kit (Zymo Research, #D5005). The converted product was 3’ tailed by dATP with terminal deoxynucleotide transferase (Enzymatics, #P7070L) in the presence of a tail-controlling adaptor, as described for the Tequila 7 N bisulfite capture sequencing assay^121^, and ligated to the adaptor with *E. coli* DNA ligase (Takara, #2161). The ligated product was amplified with a G-quadruplex-specific common primer (NGS-1290, 5’-<u>GGAGGGGAGGGGAGAGGGG</u>-3’) and an adaptor-specific common primer for 12 cycles with Phusion U (Thermo Fisher, #F555L). One-tenth of the amplification product was subjected to another round of amplification using Illumina P5/P7 primers. The amplified product was recovered by column purification and sequenced on an Illumina NovaSeq 6000 machine in PE150 format to 5-10 M reads.

### Sequencing data processing

The reference genomes used in this study were hg38 (human), hg19 (human: bisulfite sequencing only), mm10 (mouse, both routine and bisulfite sequencing), hg38-mm10 (human/mouse mixed cells for quantitative ATAC-seq/CUT&Tag), dm6 (*D. melanogaster*), sc3 (*S. cerevisiae)*, ce11 (*C. elegans*), and danRer10 (*D. rerio*). CUT&Tag, CUT&Flow, ChIP-seq, ATAC-seq, G4Access, MNase-seq, and SNS-seq data were aligned to the reference genome using bwa (0.7.17-r1188). RNA sequencing data, including total RNA-seq and TT-seq data, were aligned to the respective reference genomes using STAR (version 2.7.10a_alpha_220818). The bisulfite sequencing data were aligned to the reference genome using bwa-meth (0.2.0). CpG methylation levels were determined using Pile-O-Meth (v0.1.13-3-gca82747). Peaks in the DNA sequencing data were identified by MACS2 (2.2.7.1). The BigWig files were generated by bamCoverage (3.3.0). Coverage profiles were generated by computeMatrix (3.3.0). The mouse genome coverage of the quantitative ATAC-seq/CUT&Tag data was normalized against the reference (human genome) read content in the same library. Prior to calculating the CV in the qBG4 CUT&Tag data, to control for the influence of chromatin accessibility on BG4 CUT&Tag efficiency, the qCUT&Tag signal at a particular locus was divided by the qATAC signal at the same locus.

### Enrichment analysis

Region enrichment between a given set of genomic loci and reference loci was performed by LOLA (1.32.0). Motif enrichment in each region was performed by rGADEM (2.44.1). Fisher’s exact test was used for statistical comparisons of enrichment. The R package regioneR (1.28.0) was used for the two-region set overlap enrichment significance test.

### Estimation of the total opened fraction of G4

Single-cell and bulk ATAC sequencing data were converted into peak-x-cell count tables. Estimation of the total opened fraction of G4 loci was performed with EpiTrace (version 055441a) using the function ‘EpiTraceAge_Convergenc’. The pGQS loci derived from G4Hunter for the corresponding reference genome were used as the reference.

### Estimation of the aging coefficient

The aging coefficient was calculated by the function ‘AssociationOfPeaksToAg’ from EpiTrace. The EpiTrace-derived cell age or the known accurate timing of the sample (for the *C. elegans, S. cerevisiae, D. melanogaster, D. rerio,* and mouse pMEF data) was used for correlation against the ATAC coverage of the full peak set. Aging peaks were defined as those with a Z-scale aging coefficient >3. Aging genes were defined as genes that harbored an aging peak in their TSS. To calculate the age-related coefficient profile for a given set of genes, the positions of the ATAC-seq peaks for these genes were converted into relative positions (from the TSS to the TES: 0%-100%), and the smoothened age coefficient was calculated from the age-related coefficients of all the ATAC-seq peaks covering these genes via the ‘*gam*’ method in R (4.1.3).

### Distance between loci and the replication initiation zone

Processed OK-seq data were downloaded from https://github.com/CL-CHEN-Lab/OK-Seq^98^. Replication initiation zones (IZs) in mouse and human cells were taken from published results. The distance between the IZ and G4 loci was calculated by GenomicRanges (1.48.0).

### Repli-seq data analysis

Processed high-resolution Repli-seq data were downloaded from NCBI (GSE137764). The genomic segments were classified according to whether they overlapped (1) with a G4 ChIP-seq peak, (2) with a pGQS locus, (3) with a non-pGQS aging locus, or (4) none of above. For classwise analysis, Repli-seq signals from the 16 S-phase timepoints for all genomic segments in a particular class were averaged, and the cumulative sum of Repli-seq signals across these time points was calculated. For segmentwise analysis, Repli-seq signals from the 16 S-phase timepoints for each genomic segment were used as a “replication curve”. The curves were clustered by K-means with the R package kml (v2.4.6.1).

### H3K27me3 CUT&Flow data analysis

The H3K27me3 CUT&Flow signal covering each gene was computed by Deeptools computeMatrix (3.3.0). For each gene, the signal was averaged for each sample and normalized from lowest to highest across the samples. The pGQSneg and pGQSpos genes were separately clustered according to the cell cycle phase, exhibiting the lowest normalized H3K27me3 signal (“valley”). The H3K27me3 signal of a gene was normalized using an “H3K27me3 modification curve” and was clustered by K-means with kml. To perform joint clustering of the H3K27me3 profile and Repli-seq profiles, a DNA replication curve of the genomic segment containing the gene was generated from the Repli-seq data and combined with the H3K27me3 curve. This chimeric curve was then subjected to K-means clustering by kml.

### H3K27me3 scChIP-seq analysis

A preprocessed count table of H3K27me3 scChIP-seq data was downloaded from NCBI (GSE117309). The reference hg19 pGQS loci (G4Hunter score >=1.52) were downloaded from NCBI (GSE187007) and lifted to hg38 using easyLift (0.2.1). To estimate total H3K27me3 peaks across G4 loci for each single cell, EpiTrace was applied to the count matrix, with the lift-over hg38 pGQS locus serving as a reference. Per-G4-locus H3K27me3 ChIP-seq coverage was computed by dividing the total number of H3K27me3 reads in G4 regions by the total number of G4 loci covered. Single-cell clusters were inferred from the total count matrix using Signac (1.6.0). Ancestral clones were inferred to be the single cells in a cluster that were most evenly shared between different derived populations.

### Prediction of pGQS, G4L1, and G4 thermostability

G4L1 sequences were searched by an adapted R code following the regex search code provided by Dr. José Arturo Londoño-Vallejo as used in Lombardi *et al*., 2019^122^. The putative G4 sequence was searched for in reference genomes using G4Hunter^57^. To predict G4 thermostability, DNA sequences were extracted for each genomic locus using Biostrings (2.64.0) to generate a FASTA file input for G4Boost^77^ (https://github.com/hbusra/G4Boost, version 0de3876).

### G4 DNA methylation analysis via G4AMP and MArchPCR

Per-CpG site sequencing coverage and relative frequency of C reads (methylated C) from G4AMP or MArchPCR sequencing results were taken from the Pile-O-meth output. For linear correlation between sample age and CpG-DNA methylation level, a custom script adapted from scAge^123^ was used following the protocol described in a previous publication^7^. Maximum likelihood estimation of sample age and determination of the 95% confidence intervals from the G4AMP data were performed with the transplanted scAge^123^ function in R based on the CpG loci, which showed a correlation coefficient < −0.7 and *p* value < 0.01. To determine the TF-associated G4 DNA methylation status, TF motifs were extracted from the JASPAR2022 database (0.99.7) and matched to G4Hunter-predicted pGQS (GSE187007, lifted from mm9 to mm10) using the motifmatchr (1.18.0) function ‘matchMotifs’ with the parameters “cutoff=5e-05” and “width=7”. G4 loci matching a particular TF were collected. For each sample, a particular TF-associated DNA methylation level was calculated as the total C reads covering CpG on TF-associated loci divided by the total C+T reads covering CpG on the same set of loci. A linear correlation between TF-associated DNA methylation and sample age was calculated in R (4.1.3). The same group of non-progeroid donors is shown in Fig. 5c and Fig. 6f (for comparison to the progeroid group). The “CTRL” donors in Fig. 6d were <=15 years old from the same non-progeroid group.

### DNA methylation maintenance analysis via Hammer-seq and Repli-BS-seq

Processed stranded CpG methylation read counts of Hammer-seq data were downloaded from NCBI (GSE131098). CpGs were classified according to whether they were within the vicinity of 100 bp of the pGQS site or G4 ChIP-seq site or if they corresponded to the solo-WCGW sites in the partially methylated domain (PMD) ^9^. The daughter strand was defined as the read strand with less DNA methylation (C-reads). If two strands had similar DNA methylation levels, then a random strand was chosen as the daughter strand. For each pair of parent-daughter strands, the parental methylated loci were defined as “C” bases in the parent strand, and “C” bases in the daughter strand were considered “maintained”. The methylation maintenance level of a particular CpG class in a sample was calculated as the mean probability of maintained methylation in daughter strands for these CpGs.

To analyze methylation maintenance in Repli-BS-seq, processed bisulfite sequencing data were downloaded from NCBI (GSE82045). CpGs were classified as described for the Hammer-seq experiments. Since no strand pair information was available, the methylation maintenance level was calculated as the methylation level normalized between timepoints (0% (min) - 100% (max)).

### Transcription factor activity inference from scATAC

Transcription factor activity was computed by ChromVAR in ArchR (1.0.1) from the pMEF SHARE-seq dataset from a previous publication^7^. Single-cell ages were estimated from the same dataset using EpiTrace. A linear correlation between TF activity and single-cell age was calculated in R (4.1.3).

## Code and data availability

The codes used in this study were deposited on GitHub (link: https://github.com/MagpiePKU/G_quadruplex_aging_paper_2024). Raw mouse sequencing data and processed human sequencing data were deposited on the CNGB GSA.

## Author contributions

YZ conceived the concept and led the study. WJ, JZ, YX and LJ performed the experiments. WJ, FC, JF and YZ performed the bioinformatic analysis. KQ and HJ collected the biospecimens. YX, HJ and YZ oversee the study. WJ, YX and YZ wrote the paper, with input from all the authors.

## Acknowledgments

The authors would like to thank all the authors who shared publicly available datasets used in this study. Dr. Qi Li at the National Institute of Biological Sciences, Beijing, China kindly provided HEK293T cells for the qATAC-seq and qCUT&Tag experiments. Dr. José Arturo Londoño-Vallejo at the Curie Institute kindly provided the regex searching code for G4L1 sequences. Dr. Srinivas Ramachandran at the University of Colorado Anschutz Medical Campus kindly provided the not yet published H3K27me3 CUT&Flow sequencing data. The excellent technical assistance of Ms. Mengxue Yu and Ms. Yayun Fang at Wuhan University is gratefully acknowledged. We would also like to thank Dr. Xiaoling Li at the National Institute of Environmental Health Sciences (NIEHS) for helpful discussion and critical reading. Part of the analysis was performed on the High Performance Computing Platform of the Center for Life Science (Peking University). This study is supported by the National Natural Science Foundation of China (82273065 and 82372654). The funders played no role in the study design, data collection and analysis, decision to publish, or preparation of the manuscript.

