## Supplementary Information for "A universal molecular mechanism driving aging"

Extended Data Figures: Pages 2-20

Extended Data Tables: Pages 21-22

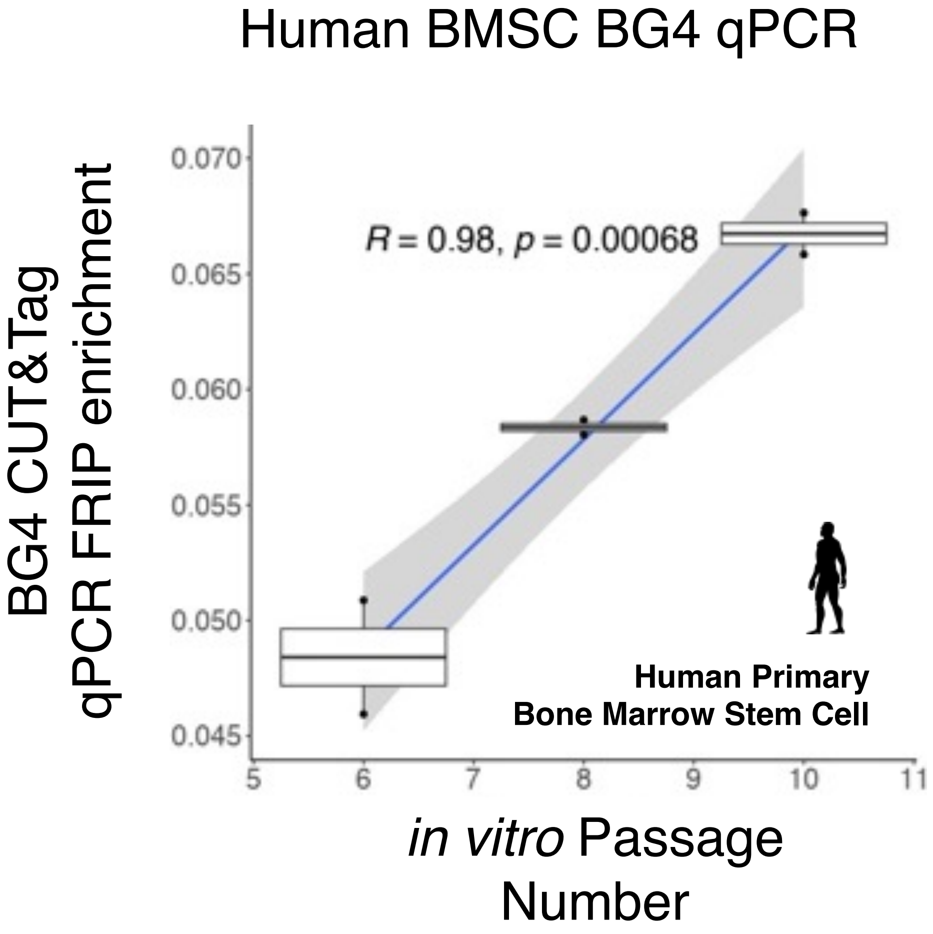


**Extended Data Figure 1. G4 qPCR enrichment against library content for human BMSCs.** qPCR amplification results with the G4 primer and Illumina P7 primer from the BG4 CUT&Tag libraries of hBMSCs are shown.

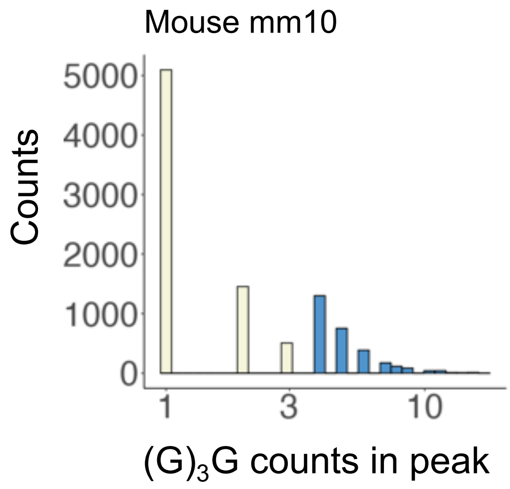


**Extended Data Figure 2. Distribution of G3/G counts in each mouse ATAC peak.** The blue columns are the peaks with >3 G3/G repeats.

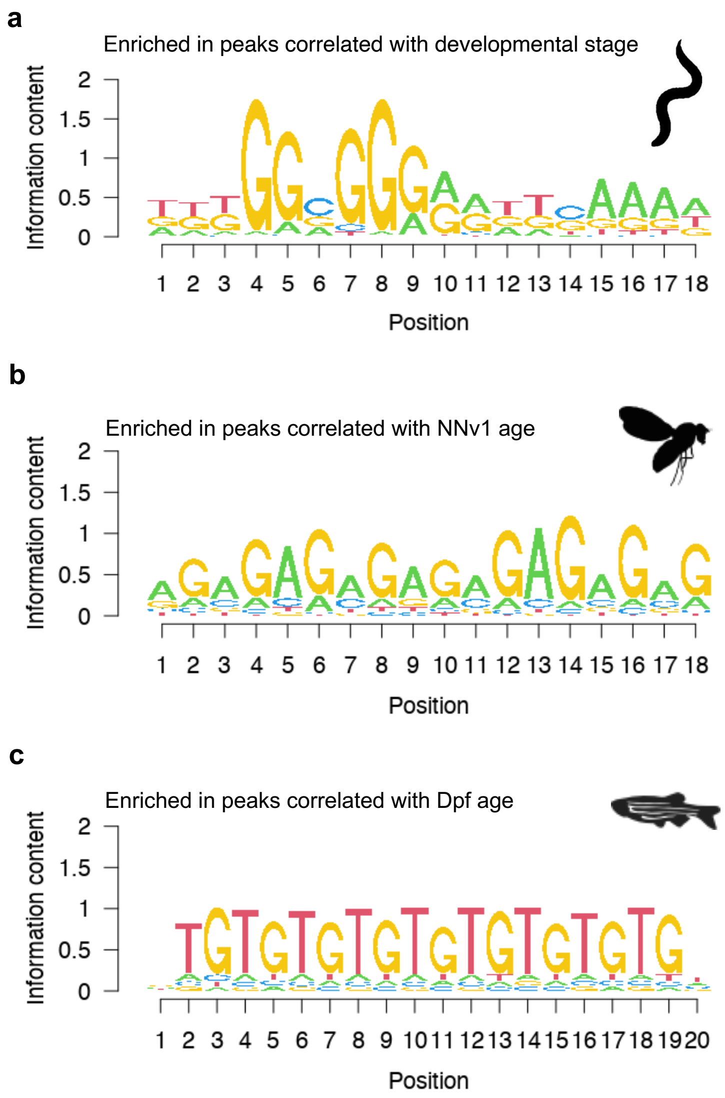


**Extended Data Figure 3. Top enriched motif associated with age-associated chromatin opening.**

**(a)** Top enriched motif in the *C. elegans* development ATAC dataset. **(b)** Top enriched motif in the *D. melanogaster* embryonic development scATAC dataset. **(c)** Top enriched motif in the *D. rerio* neural crest development scATAC dataset.

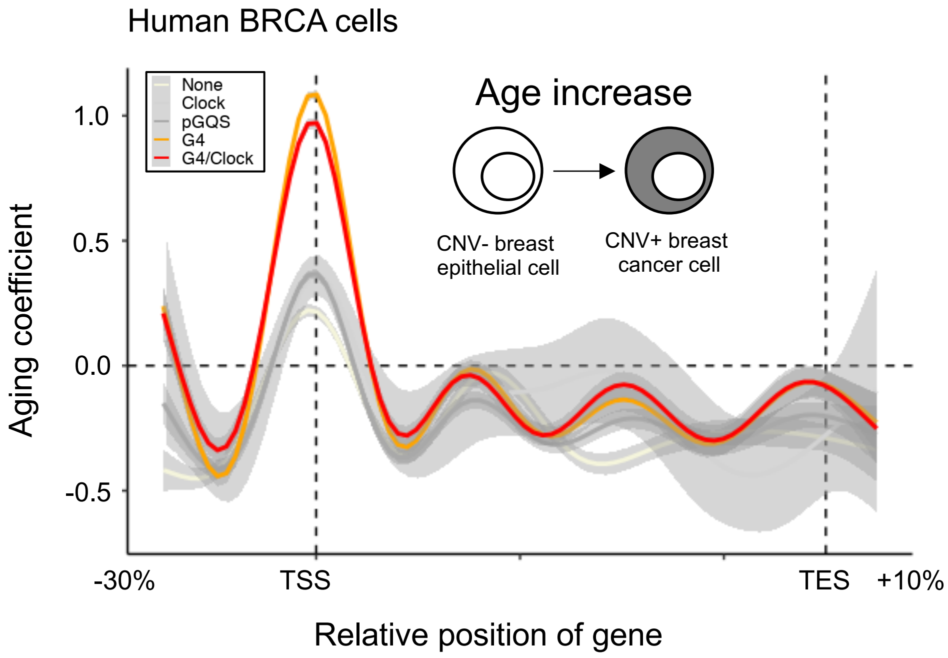


**Extended Data Figure 4. Aging coefficient profiles of genes in human breast cancer cells.** The genes were classified as (1) harboring ClockDML in the TSS; (2) harboring pGQS in the TSS; (3) harboring the G4-ChIP-seq signal in the TSS; (4) harboring both ClockDML and the G4-ChIP-signal; or (5) not harboring any of the above.

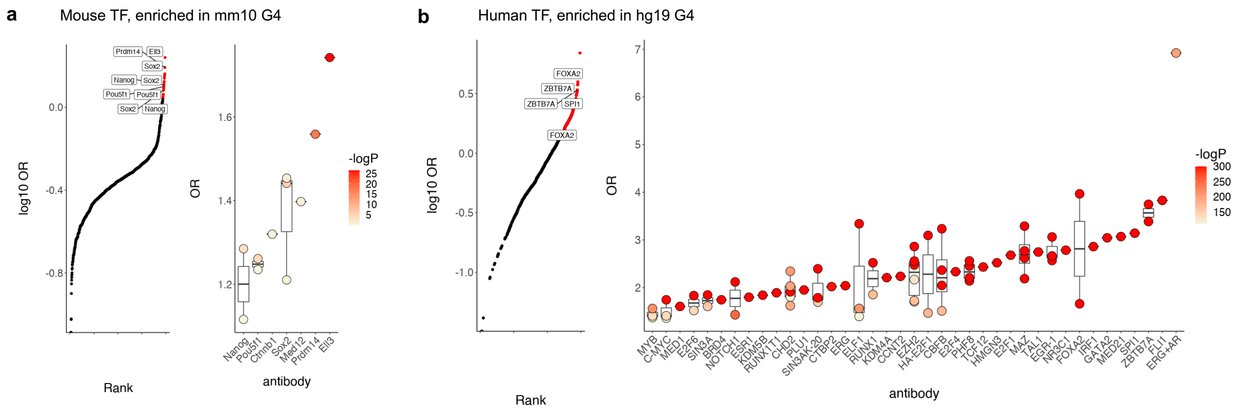


**Extended Data Figure 5. Enrichment of TF-ChIP-seq peaks at pGQS loci in the mouse and human genomes.**

**(a)** (left) Enrichment (Y-axis) vs rank (X-axis) of TF ChIP-seq peaks in pGQS in the mouse genome. The top 10% of enriched TFs are highlighted in red and labeled. (right) Enrichment (Y-axis) of the top enriched TFs. Individual ChIP-seq peak sets of the same TF are shown as dots and filled with log P-value denoting enrichment significance. **(b)** (left) Enrichment (Y-axis) vs rank (X-axis) of TF ChIP-seq peaks in pGQS in the human genome. The top 10% of enriched TFs are highlighted in red and labeled. (right) Enrichment (Y-axis) of the top enriched TFs. Individual ChIP-seq peak sets of the same TF are shown as dots and filled with log P-value denoting enrichment significance.

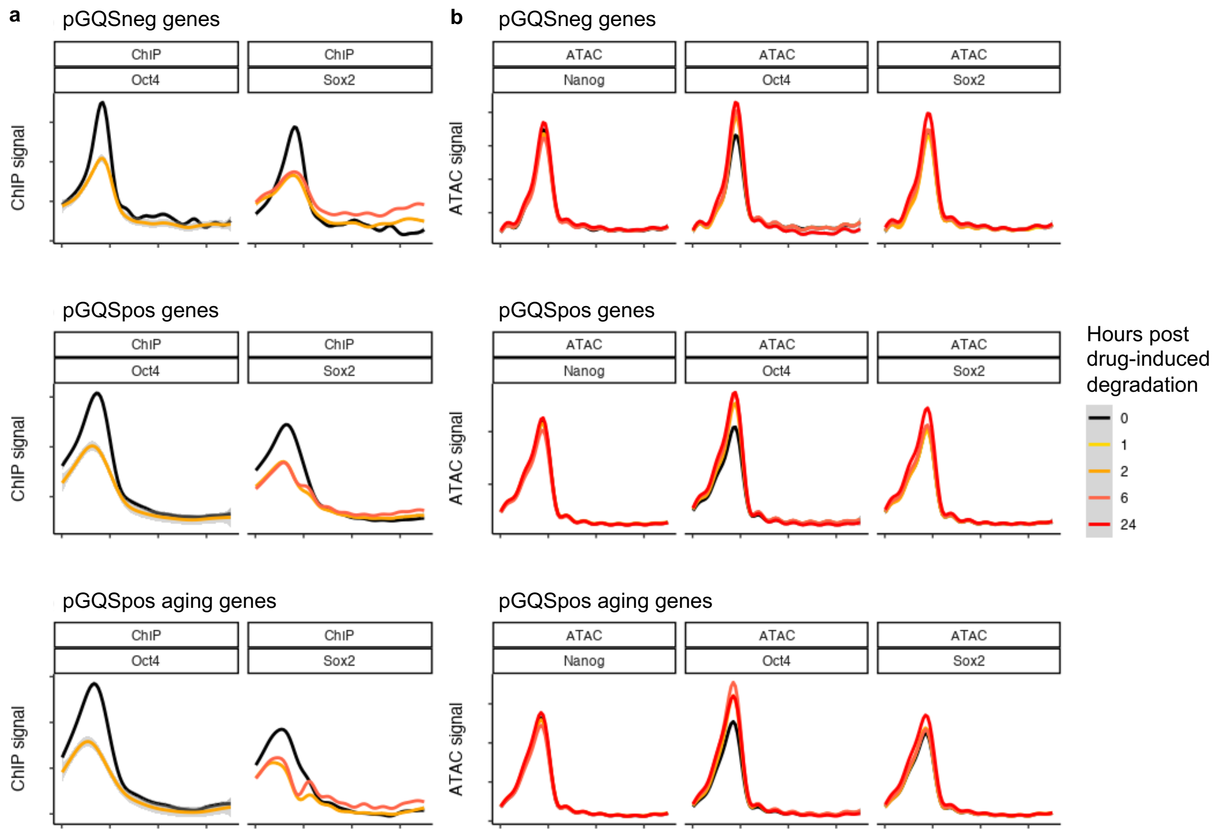


**Extended Data Figure 6. Pioneering pluripotency factors that bind to pGQS but not regulate pGQS chromatin opening.**

**(a)** ChIP-seq signal profiles of FKBP-Oct4 and FKBP-Nanog on the pGQSneg, pGQSpos, and pGQSpos aging genes. Drug-induced degradation of Oct4 and Nanog was performed for the indicated time length (denoted by color) for each sample. The control state is 0 hours. **(b)** ATAC-seq signal profiles of FKBP-Oct4, FKBP-Nanog and FKBP-Sox2 cells for the pGQSneg, pGQSpos, and pGQSpos aging genes. Drug-induced degradation of Oct4, Nanog and Sox2 was performed for the indicated time length (denoted by color) for each sample. The control state is 0 hours.

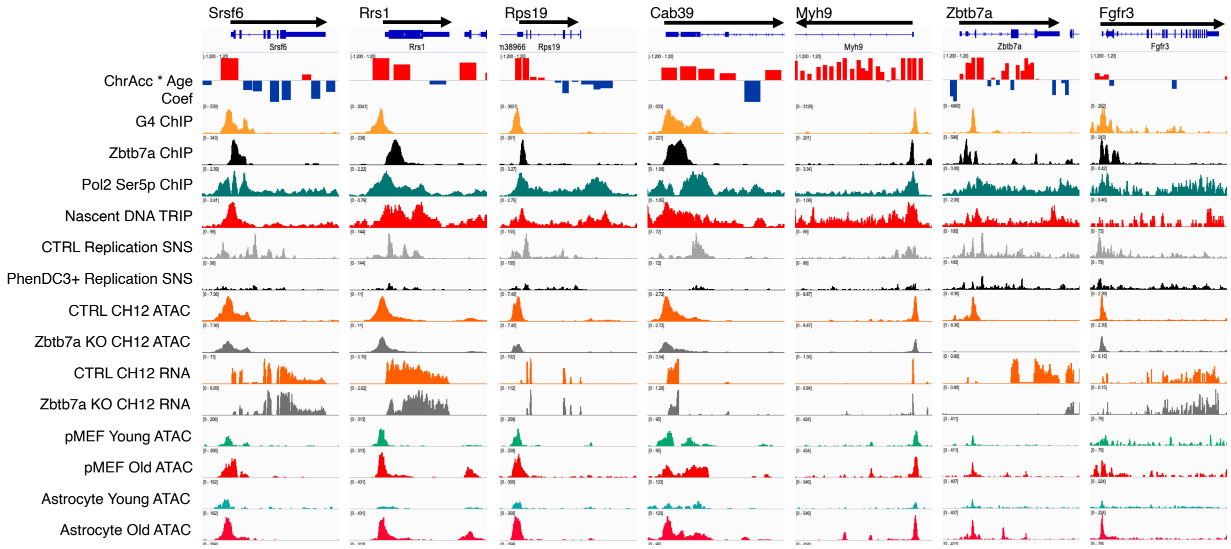


**Extended Data Figure 7. Enrichment of G4, Zbtb7a binding, RNApol2 Ser5p, nascent DNA TRI, and G4-responsive DNA replication in aging genes in the mouse genome.** The G4 ChIP-seq track was obtained from mESCs. The Zbtb ChIP-seq track was obtained from Ch12 cells. The pMEF and pAST ATAC results are also shown for comparison.

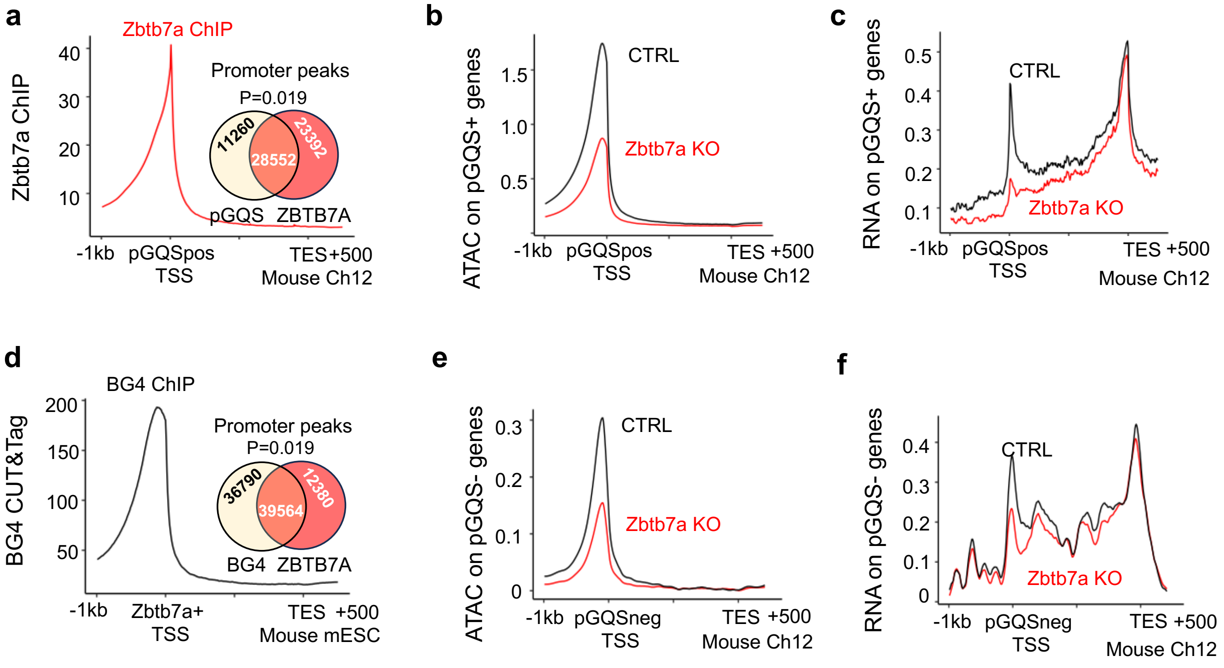


**Extended Data Figure 8. Zbtb7a regulates G4 locus chromatin.**

**(a)** ChIP-seq signal profiles of Zbtb7a on pGQSpos genes in mouse Ch12 cells. Inset: overlaps between Zbtb7a ChIP-seq peaks and pGQS loci (P=0.019 according to the permutation test). **(b)** ATAC-seq profiles of pGQSpos genes in control and Zbtb7a KO Ch12 cells. **(c)** RNA-seq profiles of pGQSpos genes in control and Zbtb7a KO Ch12 cells. **(d)** ChIP-seq signal profiles of G4s of the pGQSpos gene in mouse mESCs. Inset: overlaps between Zbtb7a and G4-ChIP-seq peaks (P=0.019 according to the permutation test). **(e)** ATAC-seq profiles of pGQSneg genes in control and Zbtb7a KO Ch12 cells. **(f)** RNAseq profiles of pGQSneg genes in control and Zbtb7a KO Ch12 cells.

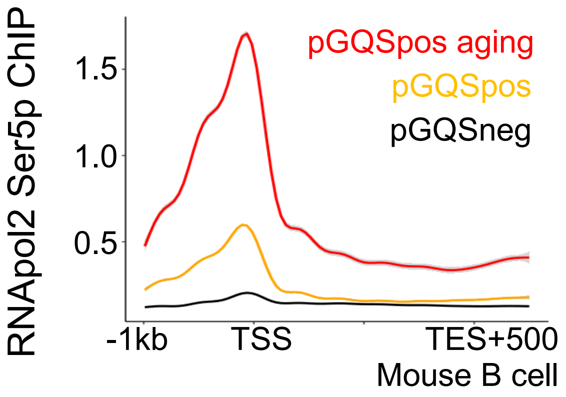


**Extended Data Figure 9. RNApol2 Ser5p ChIP-seq profile of the pGQSneg, pGQSpos and pGQSpos aging genes in mouse B cells.** Black: pGQSneg; orange: pGQSpos; and red: pGQSpos aging genes.

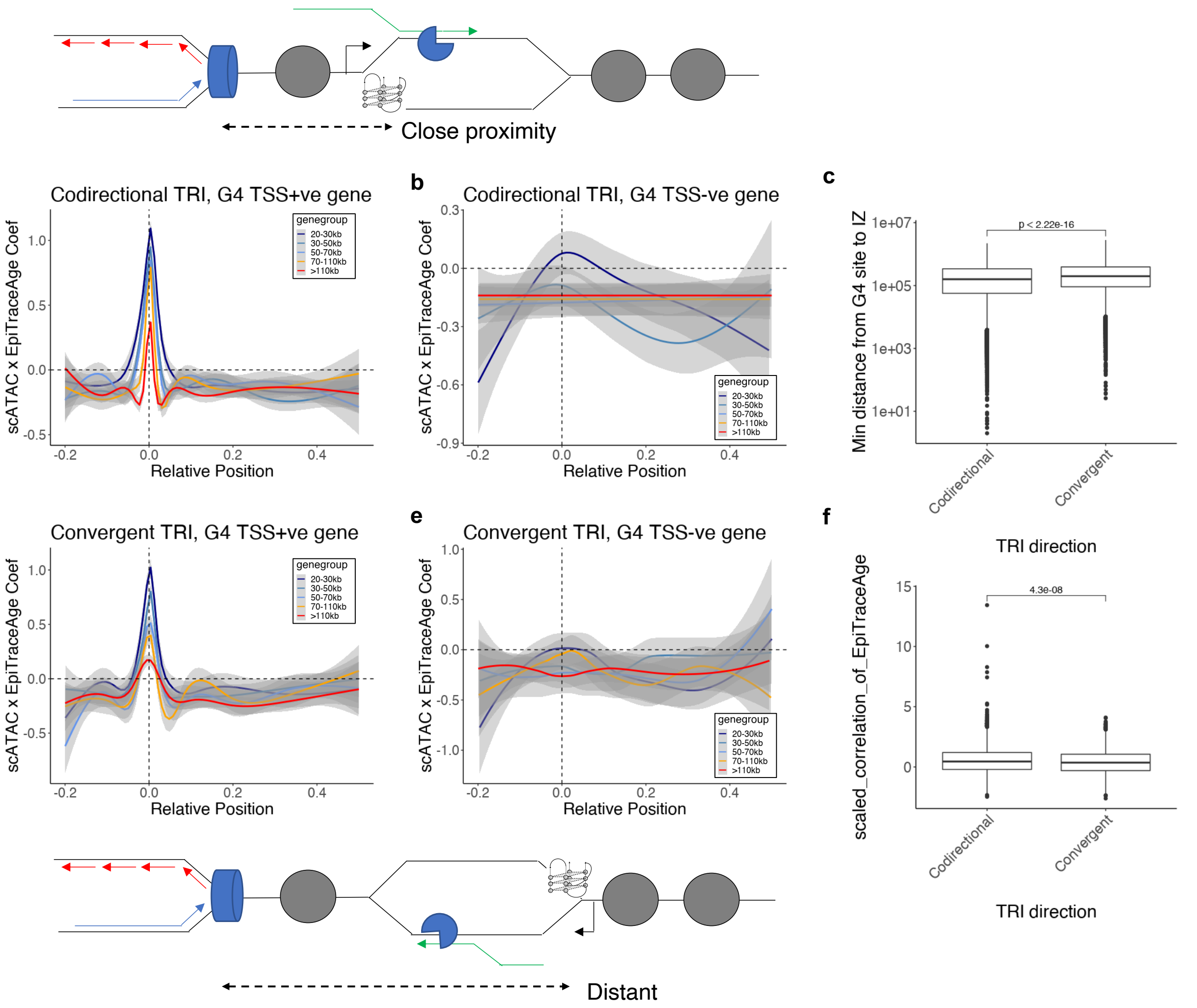


**Extended Data Figure 10. TRI influences the aging coefficient of loci.**

**(a)** Smoothened aging coefficient of pGQSpos genes with codirectional TRI. Genes were further classified according to their length. **(b)** Smoothened aging coefficient of pGQSneg genes with codirectional TRI. Genes were further classified according to their length. **(c)** Minimal distance between the pGQS locus and the replication initiation zone (IZ) in the codirectional or convergent TRI class of genes. **(d)** Smoothened aging coefficient of pGQSpos genes with convergent TRI. Genes were further classified according to their length. **(e)** Smoothened aging coefficient of pGQSneg genes with convergent TRI. Genes were further classified according to their length. **(f)** Aging coefficient of pGQS loci in the codirectional or convergent-TRI class of genes.

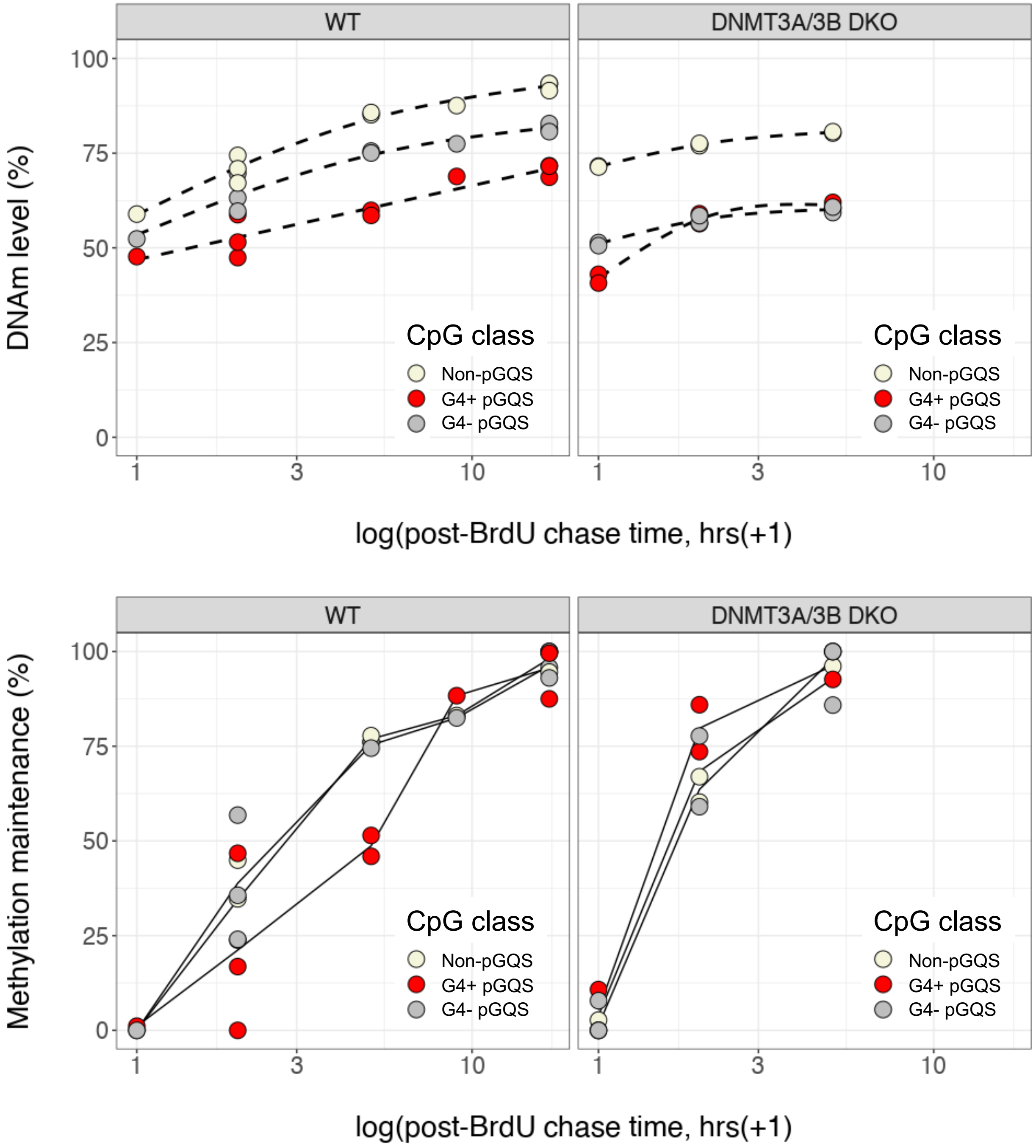


**Extended Data Figure 11. G4 impairs DNA methylation maintenance in human embryonic stem cells.**

**(a)** DNA methylation level (Y-axis) of each class of CpG in wild-type HUES64 cells pulsed with BrdU and chased with thymidine for the indicated times (X-axis). **(b)** DNA methylation level (Y-axis) of each class of CpG in DNMT3A/3B double knockout (DKO) HUES64 cells pulsed with BrdU and chased with thymidine for the indicated times (X-axis). **(c)** DNA methylation maintenance level (Y-axis), defined as the normalized (min-max) methylation level, of each class of CpG in wild-type HUES64 cells pulsed with BrdU and chased with thymidine for the indicated times (X-axis). **(b)** DNA methylation maintenance level (Y-axis) of each class of CpG in DNMT3A/3B DKO HUES64 cells pulsed with BrdU and chased with thymidine for the indicated times (X-axis).

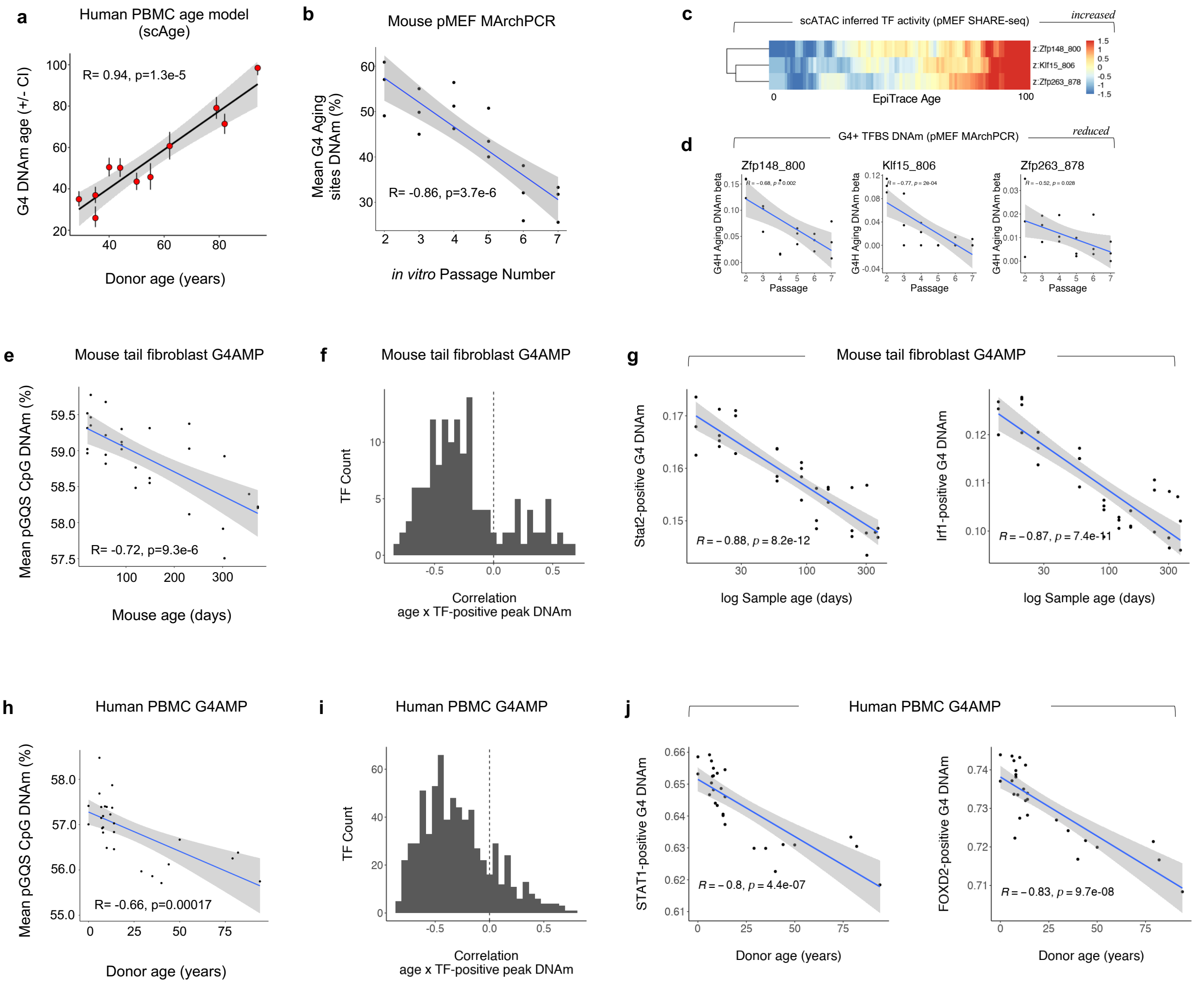


**Extended Data Figure 12. G4 DNA hypomethylation *in vivo* during aging.**

**(a)** Real (X-axis) and G4 DNAm-predicted age (scAge, Y-axis) of human PBMC samples sequenced by G4AMP. The 95% confidence intervals of the predicted biological age are also shown. **(b)** Mean DNA methylation level of age-associated CpGs in the vicinity of G4s in mouse pMEFs. **(c)** SHARE-seq scATAC data showing the transcription factor activities (color) of Zfp148, Zfp263, and Klf15 with respect to single-cell age (X-axis) in pMEFs. The inferred transcription factor activities are shown as Z-scaled normalized values. **(d)** Mean DNA methylation of G4 loci bound by Zfp148, Zfp263, and Klf15 in the same batch of pMEFs of different ages. **(e)** Mean DNA methylation of G4 loci in primary mouse tail fibroblast samples of different ages. **(f)** The distribution of correlation coefficients between the mean DNA methylation level at the TF-bound G4 locus and sample age in the mouse tail fibroblast dataset. **(g)** Mean DNA methylation of Stat2-bound and Irf1-bound G4 loci in primary mouse tail fibroblast samples of different ages. **(h)** Mean DNA methylation of G4 loci in human PBMC samples of different ages from healthy donors. **(i)** The distribution of correlation coefficients between the mean DNA methylation level of TF-bound G4 loci and sample age in the human PBMC dataset. **(j)** Mean DNA methylation of the STAT1-bound and FOXD2-bound G4 loci in the human PBMC dataset.

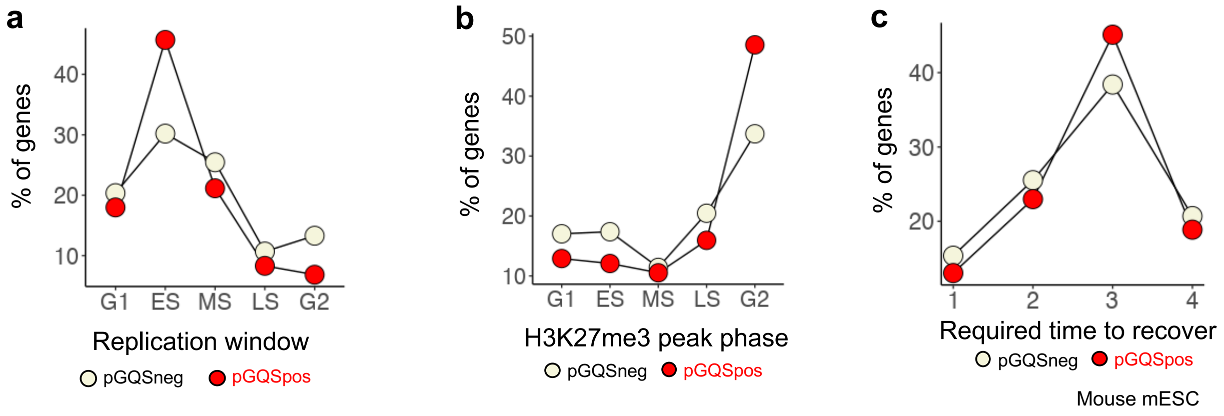


**Extended Data Figure 13. H3K27me3 recovery during the cell cycle in the pGQSpos and pGQSneg genes in mouse mESCs.**

**(a)** Percentage of genes that initiate replication (showing the lowest H3K27me3 signal) in the indicated cell cycle phase. **(b)** Percentage of genes that reached the maximal H3K27me3 signal in the indicated cell cycle phase. **(c)** Percentage of genes that required the indicated time (number of cell cycle phases, G1/ES/MS/LS/G2) to reach the maximal H3K27me3 value from the replication initiation phase.

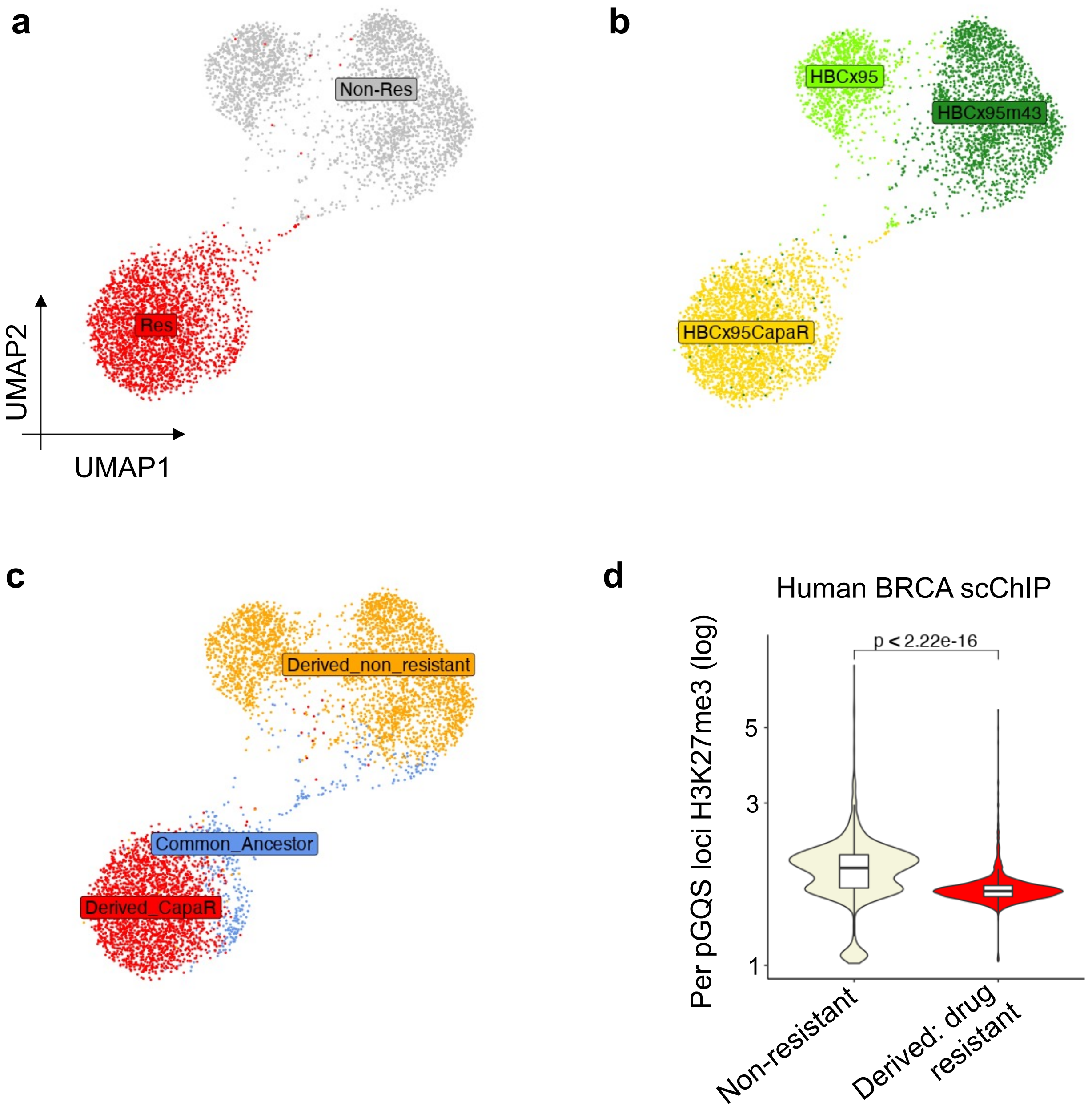


**Extended Data Figure 14. H3K27me3 at the G4 locus is reduced during human BRCA cell aging**

**(a)** UMAP plot of human BRCA cells from the scChIP-seq experiment, labeled by the capecitabine resistance state of each sample. **(b)** UMAP plot of human BRCA cells from the scChIP-seq experiment, labeled by sample name. **(c)** UMAP projection of human BRCA cells from the scChIP-seq experiment, labeled by scChIP-seq profile-based single-cell clusters. **(d)** Mean H3K27me3 reads on each pGQS locus in the non-resistant or the derived drug-resistant cells, as in (a).

**a**

| Patient | ID | Age | Mutation | Mutation class |
|---|---|---|---|---|
| P1 | 1902755 | 10 | ERCC8 Exon 4 rearrangement / Exon 4 rearrangement | Hypomorph I |
| P2 | 1902750 | 8 | ERCC8 Exon 4 rearrangement / c.394_398delTTACA | Hypomorph I |
| P3 | 1902752 | 7 | ERCC8 Exon 4 rearrangement / IVS2+5delG | Hypomorph I |
| P4 | 1902751 | 6 | ERCC8 Exon 4 rearrangement / Exon 4 rearrangement | Hypomorph I |
| P5 | 1902753 | 13 | ERCC8 Exon 4 rearrangement / Deep intronic mutation | Hypomorph II |
| P6 | 1902749 | 14 | ERCC8 Exon 4 rearrangement / Complex | Hypomorph II |
| P7 | 1902756 | 9 | ERCC8 Exon 4 rearrangement / Large fragment deletion | Null |

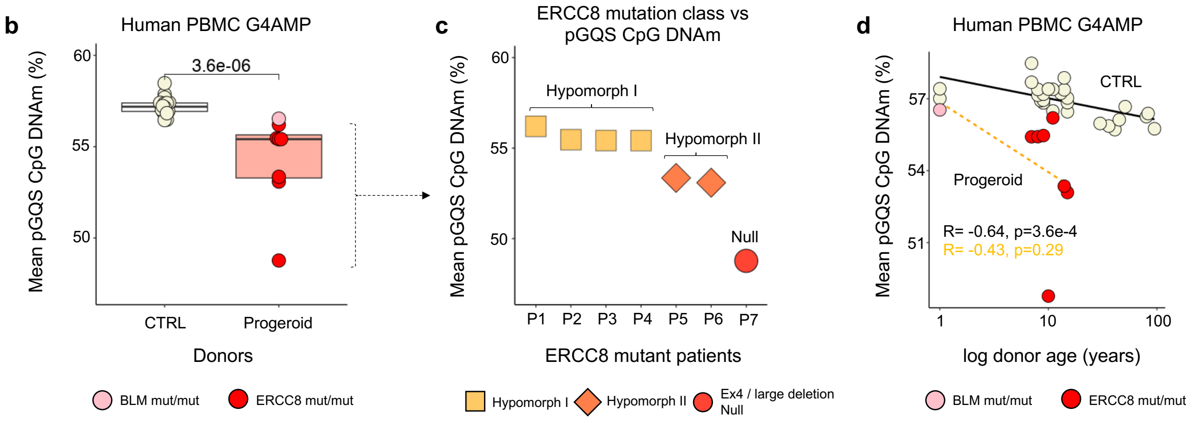


**Extended Data Figure 15. Details of the progeroid patients.**

**(a)** Genetic mutation details of each CSB patient. **(b)** Mean pGQS CpG DNAm of age-matched control (CTRL) or progeroid patients. **(c)** Mean pGQS CpG DNAm of ERCC8 mutant patients. **(d)** Relationship between age and the mean pGQS CpG DNAm in control or progeroid patients.

### Extended Data Tables S1-S2

**Extended Data Table 1. Public datasets used in this study.**

| Accession.Code / URL | Annotation |
|---|---|
| GSE131098 | Hammerseq |
| GSE82045 | Repli-BS-seq |
| https://github.com/CL-CHEN-Lab/OK-Seq | OK-seq |
| GSE165128 | Ch12 cells (Zbtb7a KO ATAC-seq, Zbtb7a KO RNA-seq) |
| GSE209527 | Sox2 Oct4 Nanog ChIP/ATAC |
| GSE178668 | G4 ligand-treated TT-seq |
| GSE52285 | WRN KO ChIP-seq |
| GSE161410 | TRIPn-seq and RNAPol2 Ser5p ChIP-seq |
| GSE137764 | High resolution Repli-seq |
| GSE126477 | SNS-Seq |
| GSE118581 | Yeast aging ATAC |
| GSE114494 | C.elegans aging ATAC |
| GSE190149 | Drosophila embryonic development scATAC |
| GSE178969 | Zebrafish neural crest development scATAC |
| GSE74912 | Human blood cell ATAC |
| GSE188461 | Human chemically induced pluripotent stem cell scATAC |
| GSE198639 | Human breast cancer scATAC |
| GSE152216 | Human breast cancer G4 ChIP-seq |
| GSE173103 | mESC G4 CUT&Tag |
| GSE187007 | G4Access |
| GSE117309 | Human breast cancer single-cell ChIP-seq (H3K27me3) |
| GSM803473 | Human ZBTB7A ChIP-seq |
| GSE122937 | Mouse Zbtb7a ChIP-seq |
| GSE209818 | Mouse mESC H3K27me3 CUT&Flow |

**Extended Data Table S2. Patient information.**

| Patient | ID | Age | Mutation | Mutation class |
|---|---|---|---|---|
| ERCC8-P1 | 1902755 | 10 | ERCC8 Exon 4 rearrangement / Exon 4 rearrangement | Hypomorph I |
| ERCC8-P2 | 1902750 | 8 | ERCC8 Exon 4 rearrangement / c.394_398delTTACA | Hypomorph I |
| ERCC8-P3 | 1902752 | 7 | ERCC8 Exon 4 rearrangement / IVS2+5delG | Hypomorph I |
| ERCC8-P4 | 1902751 | 6 | ERCC8 Exon 4 rearrangement / Exon 4 rearrangement | Hypomorph I |
| ERCC8-P5 | 1902753 | 13 | ERCC8 Exon 4 rearrangement / Deep intronic mutation | Hypomorph II |
| ERCC8-P6 | 1902749 | 14 | ERCC8 Exon 4 rearrangement / Complex | Hypomorph II |
| ERCC8-P7 | 1902756 | 9 | ERCC8 Exon 4 rearrangement / Large fragment deletion | Null |
| BLM-P1 | 2102456 | 0 | BLM c.1544dup / c.3300A>G | PAT/LP |
